# Replication-induced DNA secondary structures drive fork uncoupling and breakage

**DOI:** 10.1101/2022.11.18.517070

**Authors:** Sophie L. Williams, Corella S. Casas-Delucchi, Federica Raguseo, Dilek Guneri, Yunxuan Li, Masashi Minamino, Emma Elisabeth Fletcher, Joseph T. P. Yeeles, Ulrich F. Keyser, Zoë A. E. Waller, Marco Di Antonio, Gideon Coster

**Affiliations:** Genome Replication lab, Division of Cancer Biology, Institute of Cancer Research, Chester Beatty Laboratories, 237 Fulham Road, London SW3 6JB, UK; Imperial College London, Chemistry Department, MSRH, 82 Wood Lane, London W12 0BZ, UK; Institute of Chemical Biology, MSRH, 82 Wood Lane, London W12 0BZ, UK; UCL, School of Pharmacy, 29-39 Brunswick Square, London WC1N 1AX, UK; Cavendish Laboratory, University of Cambridge, Cambridge, UK; Francis Crick Institute, 1 Midland Rd, London NW1 1AT, UK; MRC Laboratory of Molecular Biology, Cambridge, UK

## Abstract

Sequences that can form DNA secondary structures, such as G-quadruplexes (G4s) and intercalated-Motifs (iMs), are abundant in the human genome and play a range of physiological roles. However, they can also pose a challenge to the replication machinery and in turn threaten genome stability. Multiple lines of evidence suggest G4s interfere with replication, but the underlying mechanism remains unclear. Moreover, there is a lack of evidence of how iMs affect the replisome. Here, we reconstitute replication of physiologically derived structure-forming sequences to find that a single G4 or iM is sufficient to arrest DNA replication. Direct single molecule structure detection within solid-state nanopores reveals structures form as a consequence of replication. A combination of genetic and biophysical characterisation establishes that structure forming capacity is a key determinant of replisome arrest. Mechanistically, replication fork arrest is caused by impaired synthesis, resulting in helicase-polymerase uncoupling. Significantly, iMs also induce breakage of nascent DNA. Finally, stalled forks are only rescued by a specialised helicase, Pif1, but not Sgs1 or Chl1. Altogether, this study provides a potential mechanism for quadruplex structure formation and resolution during replication and highlights G4s and iMs as endogenous sources of replication stress, which may explain their genomic instability and mutation frequencies in cancer.

## Introduction

Eukaryotic DNA replication is a highly regulated process carried out by a complex molecular machine known as the replisome (Bell and Labib, 2016). Parental duplex DNA is unwound by the replicative helicase, CMG, and nascent DNA is subsequently synthesised by polymerase ε on the leading strand, or polymerase δ on the lagging strand. The replisome must replicate all regions of the genome accurately while encountering various obstacles such as DNA damage, protein barriers and transcription-replication collisions (reviewed in (Zeman and Cimprich, 2014)). All of these can lead to replication stress, which poses a challenge to genome integrity (Gaillard et al., 2015).

Another potential barrier to replisome progression is the DNA template itself. Aside from the canonical B-DNA conformation, certain DNA sequences can fold into secondary structures, particularly from ssDNA exposed during replication. Examples of well-characterised secondary structures include hairpin (Gacy et al., 1995, Nadel et al., 1995), G-quadruplex (G4) (Fry and Loeb, 1994) and intercalated-Motif (iM) (Gehring et al., 1993) structures, which are thought to act as barriers to replication. For example, hairpin-forming repeats cause replication stalling in yeast (Voineagu et al., 2008). Moreover, two unbiased studies have mapped sites of replication fork collapse *in vivo*, highlighting poly(dA) sites (Tubbs et al., 2018) and a variety of structure-forming repeats (Shastri et al., 2018). Therefore, secondary structures may be responsible for the fact that repetitive sequences are unstable and give rise to genomic instability (reviewed in (Brown and Freudenreich, 2021)).

G4 structures are formed from guanine-rich sequences through the stacking of G quartets formed by Hoogsteen base pairing (Sen and Gilbert, 1988). Their abundance increases during S-phase, suggesting they arise as a consequence of replication (Biffi et al., 2013, Di Antonio et al., 2020). G4-forming sequences are defined by a motif consisting of four consecutive tracts of at least three guanines (G-tracts), separated by one to seven non-G nucleotides (G_3_N_1-7_G_3_N_1-7_G_3_N_1-7_G_3_) (Huppert and Balasubramanian, 2005) which are referred to as ‘loops’. The length and composition of these loops affects the stability of the G4 structure. For example, shorter loops composed of thymine residues correlate with more thermally stable structures, while longer loops result in less stable structures (Piazza et al., 2015).

G4s can exist in a variety of structural topologies, depending on the orientation of the G-rich strands relative to one another (parallel, anti-parallel or hybrid (Ou et al., 2008)). G4-forming sequences are abundant in the human genome and are known to have a range of physiological roles. Firstly, they have been associated with telomere maintenance (Maiti, 2010). Secondly, they are enriched in promoter regions (Chambers et al., 2015) where they can affect the transcriptional state of genes (reviewed in (Robinson et al., 2021)). More recently, ChIP-Seq of G4-structures on a genome-wide scale revealed that these structures mark sites of actively transcribed genes (Hansel-Hertsch et al., 2016). Finally, they are implicated in human diseases. For example, G4s correlate with telomere fragility and induce both genetic and epigenetic instability (Papadopoulou et al., 2015, Schiavone et al., 2014, Vannier et al., 2013) and have been shown to disrupt repressive chromatin structures (Sarkies et al., 2010). Moreover, G4s, such as those found at the *c-myc* gene promoter, are enriched at mutation hotspots in an array of cancers (Wang and Vasquez, 2017, De and Michor, 2011).

i-Motifs (iMs) arise from cytosine-rich DNA and therefore, in genomic contexts, can often be found on the opposite strand to G4s, for example at telomeres (Zeraati et al., 2018). These structures are held together by hemi-protonated cytosine-cytosine base pairs (C:C+) and are stabilised by low pH (Gehring et al., 1993). Although the existence of iMs has long been established, there has been speculation about their biological relevance due to their apparent requirement for acidic conditions. However, recent studies have shown that certain iMs can form at neutral pH (Wright et al., 2017), requiring longer tracts of cytosines with a proposed consensus of four consecutive tracts of at least five cytosines, separated by one to nineteen non-C nucleotides (C_5_(N_1-19_C_5_)_3_) (Wright et al., 2017). The thermal stability of iMs can be measured similarly to G4s by calculating the melting temperature of the structure. In addition to temperature, iM stability can be affected by the local pH, often described by the transitional pH (pH_T_) - the pH at which the sequence is 50% folded (Wright et al., 2017). Similar to G4s, a longer tract of Cs and a shorter loop length creates a more stable iM. In contrast to G4s, iMs can only adopt an anti-parallel conformation. iMs can be found at transcription start sites, and in the promoters of oncogenes such as *BCL2* (Kendrick et al., 2014), *HRAS* (Miglietta et al., 2015), *KRAS* (Kaiser et al., 2017)*, c-MYC* (Simonsson et al., 2000) and *VEGF* (Guo et al., 2008).

Many studies have indicated that G4s can impact replication *in vivo.* In the absence of specialised helicases, such as Pif1, G4s positioned on the leading strand template can undergo rearrangements and mutations (Lopes et al., 2011). While the replication machinery stalls at G4-forming sequences in a variety of organisms, there is conflicting evidence of stalling on the leading versus lagging strand template (Lopes et al., 2011, Sarkies et al., 2010, Paeschke et al., 2011, Dahan et al., 2018). In addition, a recent super-resolution microscopy study demonstrated that replication-associated G4s interfere with replication stress signalling (Lee et al., 2021). Despite the abundance of evidence that G4s can interfere with replication, the underlying trigger remains unknown. Moreover, the mechanism of stalling and which replisome components are affected remain unclear.

Primer extension assays have demonstrated that structure-forming sequences inhibit synthesis. A variety of G4-forming sequences are sufficient to inhibit a range of polymerases *in vitro,* including the replicative polymerases ε and δ (Edwards et al., 2014, Lormand et al., 2013, Murat et al., 2020, Sparks et al., 2019b). There is some evidence that polymerase inhibition is affected not only by the stability of the G4, but also its topology (Takahashi et al., 2017). DNA synthesis through G4s can be aided by accessory factors such as specialised G4-unwinding helicases, including Pif1, REV1 and FANCJ (Fan et al., 2009, Ray et al., 2013, Salas et al., 2006, Sparks et al., 2019b). However, these studies were carried out with pre-folded G4 structures using isolated polymerases. Whether such structures can form in the context of dsDNA unwound by CMG and how G4s affect the complete eukaryotic replisome is unknown.

Much less is known about the effect of iM structures on DNA replication. The only available data is from primer extension assays, which have demonstrated that iMs can inhibit DNA synthesis *in vitro* (Catasti et al., 1997, Murat et al., 2020, Takahashi et al., 2017).

Recent work from our lab demonstrated that a variety of repetitive sequences stall the eukaryotic replisome. These include tracts of poly(dG)_n_ and poly(dC)_n_ sequences, which can form G4s and iMs, respectively (Casas-Delucchi et al., 2022). These results are supported by recent work from the Remus lab, which demonstrated that poly(dG)_n_ tracts stall the eukaryotic replisome when initially stabilised by an R-loop. This study also highlighted the ability of a pre-formed G4 (generated from the synthetic sequence (GGGT)_4_) to inhibit unwinding by the CMG helicase (Kumar et al., 2021). Poly(dG)_n_ and poly(dC)_n_ sequences are unique types of G4 and iM-forming sequences due to their homopolymeric nature, and the stability and type of structures they form is highly polymorphic and thus difficult to study. As such, these sequences may not be representative of physiological G4 or iM-forming sequences. Yeast telomeric DNA alone does not affect replisome progression *in vitro*, and replication stalling is only seen when the telomeric binding protein Rap1 is present (Douglas and Diffley, 2021). However, yeast telomeres are relatively weak G4-forming sequences. Whether G4 and iM-forming sequences found in the human genome are sufficient to stall replication remains unclear.

Here, we carry out an extensive study on a variety of physiologically derived G4 and iM sequences and investigate how they impact eukaryotic replication. Using a reconstituted budding yeast DNA replication system, we find that a single G4 or iM-forming sequence is sufficient to cause replisome stalling. The ability of these sequences to stall replication correlated with their ability to form stable structures, strongly suggesting that secondary structure formation was the underlying trigger for replication stalling. Interestingly, CMG was able to unwind past these sequences, while DNA synthesis by polymerase ε was inhibited, leading to helicase-polymerase uncoupling and exposure of ssDNA. Direct detection of secondary structures by single-molecule sensing using solid-state nanopores established the lack of pre-formed structures prior to replication, and conditions that enhanced replication fork uncoupling led to increased fork stalling. Together, these observations support a model whereby replication-dependent structures arise behind CMG on ssDNA exposed during unwinding, resulting in inhibition of polymerase activity. Remarkably, stalling could only be rescued by the G4-unwinding helicase Pif1, but not Sgs1 or Chl1, highlighting the specificity of this enzyme. Moreover, we found that iMs have the ability to cause DNA breakage.

Ultimately, this study describes the response of the eukaryotic replisome to a variety of physiological structure-forming motifs and highlights their ability to induce replication stress. This mechanism may provide a potential explanation as to why these sequences are both genetically and epigenetically unstable and may explain their high mutational frequencies in cancer.

## Results

### The replisome stalls at a single quadruplex-forming sequence

To establish how quadruplex-forming sequences affect the eukaryotic replisome, we cloned several well-characterised G4- and iM-forming sequences into a 9.8 kb plasmid. These were used to generate substrates for *in vitro* eukaryotic replication using purified budding yeast proteins (Yeeles et al., 2015, Yeeles et al., 2017). The G4-forming sequences tested include one of the most well-characterised G4s found in the human genome, *c-MYC* Pu22. This is a 22-nt-long segment derived from the human *c-MYC* promoter region (Dai et al., 2011) which is frequently mutated in cancers. Other G4-forming sequences tested include (i) the human telomeric repeat sequence (TTAGGG)_4_, (ii) the Bu1 +3.5 G4 motif (derived from the avian DT40 genome) which induces replication-dependent epigenetic silencing (Schiavone et al., 2014), (iii) the CEB25 L111(T) motif derived from the naturally occurring human CEB25 mini-satellite, where all three loops have been modified to a single thymine, leading to a thermally stable G4 structure (Piazza et al., 2015), (iv) the GGGGCC repeat of the C9orf72 gene which is associated with familial amyotrophic lateral sclerosis (Thys and Wang, 2015), (v) a synthetic sequence (GGGT)_4_, and (vi) a tract of poly(dG)_16_ which has previously been shown to induce replication stalling (Casas-Delucchi et al., 2022) (Table 1). Selection of iM sequences was based on their ability to form secondary structures at physiological pH (Wright et al., 2017). These exist in the human genome and are derived from the promoter regions of (i) DAP, (ii) DUX4L22, (iii) SNORD112, (iv) AC017019.1, (v) PIM1 and (vi) ZBTB7B (Table 2).

Sequences were cloned 3 kb downstream of the ARS306 origin, from which site-specific replisome loading and firing occurs (**Fig. 1A**). As replication initiates from a defined site in the template, we can infer the identity of the leading and lagging nascent strands. The structure-forming sequences we refer to throughout the manuscript are positioned on the leading strand template. The templates were linearised by digestion with AhdI before replication to avoid confounding effects of converging replisomes on circular replication templates. Upon replication initiation on the parental control substrate, two replication forks proceed from the origin in either direction, generating one longer leading strand product of 8.2kb, and one shorter leading strand product of 1.5kb. However, if the replisome stalls at a structure-forming sequence, a 3kb band appears at the expense of the longer 8.2 kb leading strand product (**Fig. S1**). Lagging strand maturation factors have been omitted in these experiments. Therefore, lagging strand products remain as Okazaki fragments and run as a smear on denaturing agarose gels (**Fig. 1B**, lane 1).

**Figure 1.**
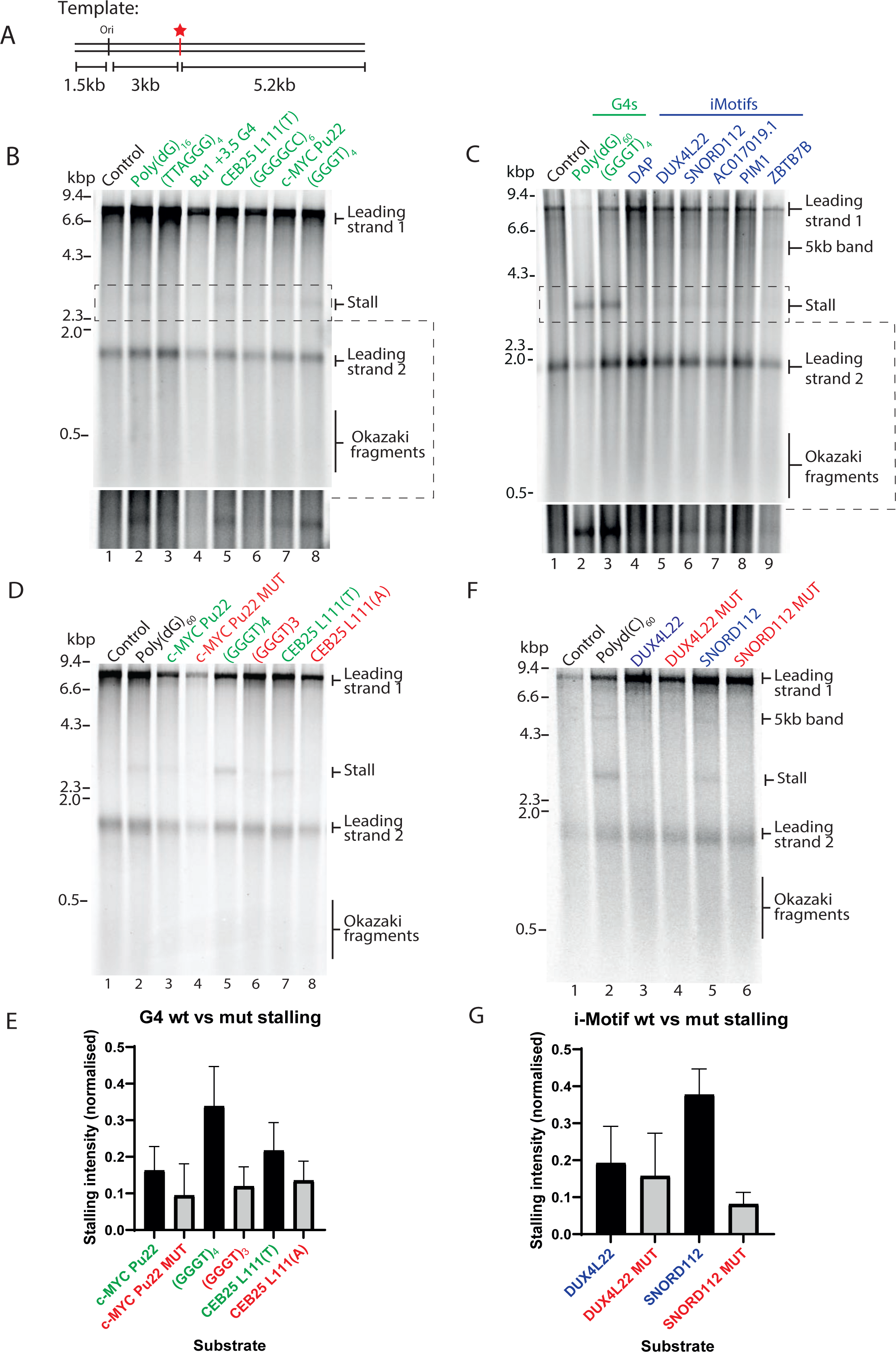
A single G4- or iM structure stalls the replisome. **(A)** Schematic of replication templates used in this study. Position of the origin of replication is marked as ‘ori’ from which replication initiates. The multiple cloning site (indicated by a red star) is positioned 3kb downstream of the origin and was used to insert various G4 or iM-forming sequences into the template. **(B)** *In vitro* replication reactions using linear substrates containing a single G4-forming sequence. Products were analysed by denaturing gel electrophoresis. Bottom panel shows increased contrast of the region containing the 3kb stall band indicated by the dashed box to better visualize stall products. **(C)** Analysis of products of *in vitro* replication reactions on linear substrates containing a G4-forming sequence (indicated in green) or a single iM-forming sequence (indicated in blue). Products were analysed on denaturing agarose gels. Bottom panel shows increased contrast of the region containing the 3kb stall band indicated by the dashed box to better visualize stall products. **(D)** Replication reactions of linear substrates containing *wildtype* or mutated G4 sequences depicted in Fig. S3A. Wildtype sequences are indicated in green and mutated sequences are indicated in red. Products are visualised on a denaturing agarose gel. **(E)** Stalling intensities of substrates as tested in (D) from three independent experiments. Stalling intensities are normalised to the intensity of ‘leading strand 2’ in each lane, and the mean ± standard error is plotted. **(F)** Analysis of replication products of linear *wildtype* iM substrates or mutated versions as shown in Fig. S3B on a denaturing agarose gel. Wildtype sequences are indicated in blue and mutated sequences are indicated in red. **(G)** Quantifications of stalling intensities induced by substrates as depicted in (F) from three independent experiments. 3kb stall band intensities were normalised to the intensity of the 1.5kb ‘leading strand 2’ product to account for variation in reaction efficiencies. Mean is plotted and error bars represent standard error of the mean.

As expected, the replicated control template generated two major leading strand products (8.2kb and 1.5kb) and Okazaki fragments (**Fig. 1B**, lane 1). In contrast, a faint but reproducible 3kb band appeared in the presence of some of the G4-forming sequences tested, indicating replisome stalling (**Fig. 1B**, lanes 2-8). The intensity of the 3kb stall band is quantified from three independent experiments in **Figure S2A**. A thermal difference spectrum (TDS) was performed for each sequence to observe the characteristic G4-profile (data not shown). The thermal stability of the G4s was assessed via UV-vis melting which revealed a positive correlation between melting temperature (T_m_) and stalling intensity, with a Pearson correlation r value of 0.76 (**Fig. S2B**). This is highlighted in Table 1, which depicts the melting temperatures in order of increasing thermal stability. Some of the G4s we tested, such as (GGGGCC)_6,_ could not be assigned a melting temperature due to aggregation issues, likely due to formation of multimolecular structures. Similarly, poly(dG)_16_ could not be accurately characterised. These sequences were therefore excluded from further studies.

Replication stalling was also reproducibly induced by a range of physiologically derived single iM sequences, although to a lesser degree than G4s (**Fig. 1C**, lanes 4-9 and **Fig. S2C**). In contrast to G4s, the intensity of replication stalling by iMs did not correlate with their relative thermal stability (**Fig. S2D**). However, there was a weak positive correlation between iM stalling and their transitional pH (pH_T_) (**Fig. S2E**, Table 2). Intriguingly, upon replication of iM sequences, a novel 5kb replication product accumulated (**Fig. 1C**, lanes 4-9). Although this product was sometimes weakly visible with G4-forming sequences, it was consistently prominent upon replication of iMs. This product corresponds to the length of leading strand 1 from the site of the structure-forming sequence downstream to the end of the template (**Fig. S1**). This could either be a result of intrinsic repriming events, or DNA breakage at the site of the iM. We investigate these possibilities later. Interestingly, the intensity of stalling seen at a single G4-forming sequence was stronger than at a single iM-forming sequence (**Fig. 1C**, compare lane 3 to lanes 4-9). We propose this is due to the lower melting temperature of iMs at physiological pH relative to G4s (Tables 1 and 2), and/or a result of different folding dynamics.

### Stalling is dependent on structure formation

To test the hypothesis that stalling is due to structure formation, we introduced mutations that abrogate or disrupt structures (**Fig. S3A, S3B**). These included removing a G-tract from (GGGT)_4_ to (GGGT)_3_ and mutating the central guanine to an adenine in each G-tract of *c-MYC* Pu22. As an intermediate experiment, we mutated the loop regions in CEB25 L111(T) from thymine to adenine residues, which still allows for G4 formation but reduces its thermal stability (Piazza et al., 2015). Consistent with our hypothesis, stalling was reduced with the mutated sequences (**Fig. 1D, 1E**). The reduction in stalling was less significant for CEB25 L111(A), likely due to the fact it is still able to form a structure, albeit weaker. Importantly, biophysical characterisation demonstrated that these sequences either form weak structures, or none at all (Table 1). As expected, the melting temperatures of the mutated G4 sequences were reduced relative to the wildtype sequences (**Fig. S3C**), resulting in a positive correlation between stalling and thermal stability (**Fig. S3E**, Pearson correlation r value of 0.7). Although the melting temperature for CEB25 L111(A) remained relatively high as it was still able to form a G4, the stalling intensity was reduced when compared to CEB25 L111(T) which is consistent with its weakened thermal stability (**Fig. 1D** and **Fig. S3C**). When the iM-forming sequences DUX4L22 and SNORD112 were disrupted for their structure-forming ability (**Fig. S3B**), stalling was reduced (**Fig. 1F, 1G**). This was less evident for DUX4L22, which is reflected in the fact that the mutations have a greater effect on the transitional pH and thermal stability of SNORD112, with a small effect on DUX4L22 (**Fig. S3D, S3F, S3G** and Table 2).

### Replisome stalling is dependent on the probability of structure formation

Stalling induced by a single G4- or iM-forming sequence was persistent, although relatively weak when compared to other types of structure-prone repeats (Casas-Delucchi et al., 2022). Larger arrays of consecutive quadruplex-forming sequences may exacerbate replisome stalling. To test this, we replicated templates containing an increasing number of consecutive G4 or iM-forming sequences (**Fig. 2A, 2B**) and found that stalling increased with the number of structure-forming sequences (quantified from three independent experiments in **Figs. S4A** and **S4B**).

**Figure 2.**
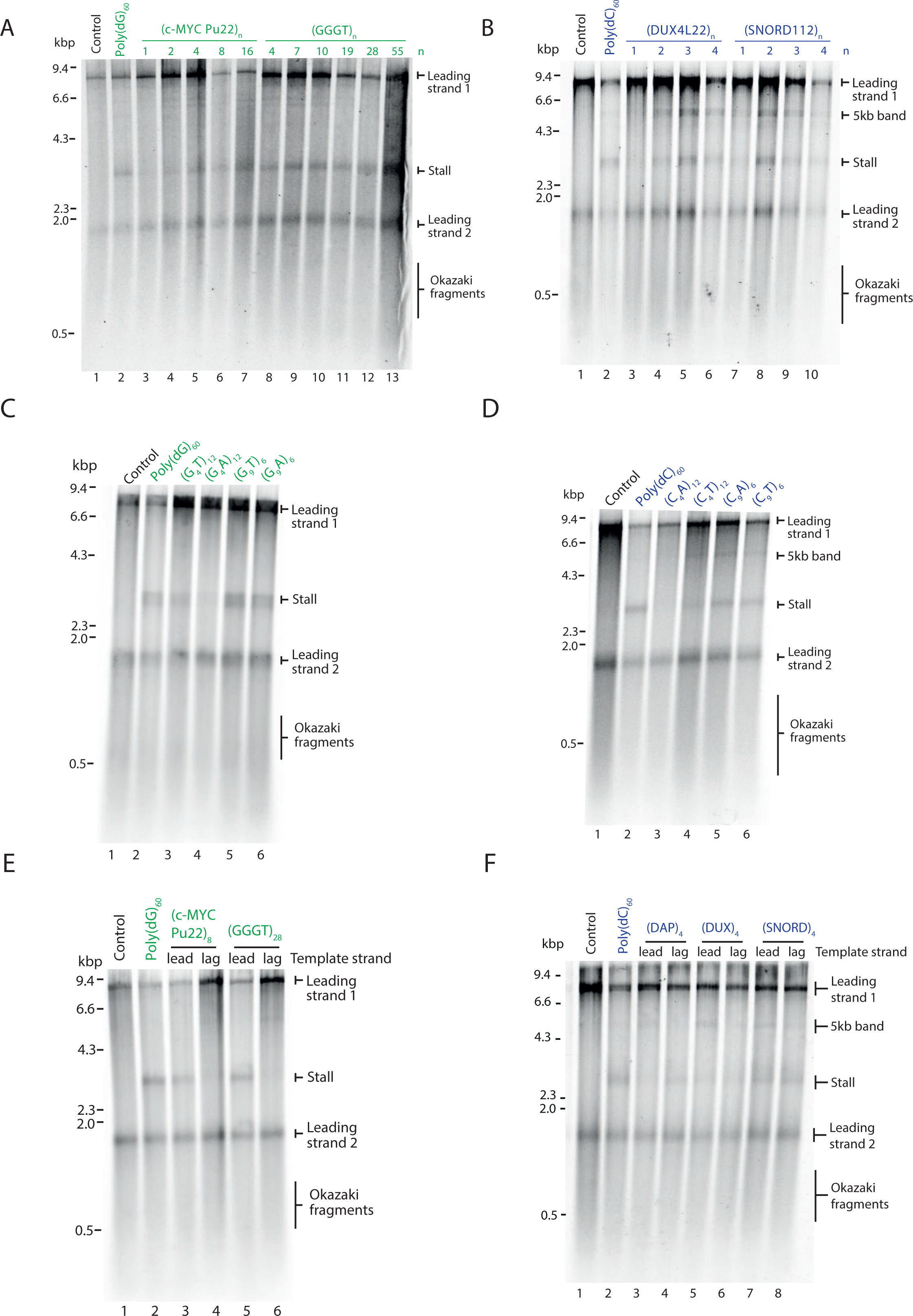
Replisome stalling is dependent on the opportunity for structure-formation and is affected by orientation. **(A)** Replication of G4 substrates containing poly(dG)_60_ or an increasing number of consecutive (*c-MYC* Pu22)_n_ or (GGGT)_n_ repeats. Products have been analysed on a denaturing agarose gel. **(B)** Replication of iM substrates containing poly(dC)_60_ or an increasing number of consecutive (DUX4L22)_n_ or (SNORD112)_n_ repeats. Products have been analysed on a denaturing agarose gel. **(C)** Substrates containing a tract of poly(dG)_60,_ or poly(dG)_60_ interrupted every 5^th^ guanine with either a thymine (G_4_T)_12_ or adenine (G_4_A)_12,_ or poly(dG)_60_ interrupted every 10^th^ guanine either a thymine (G_9_T)_6_ or adenine (G_9_A)_6_ were replicated. Replication products were analysed on a denaturing agarose gel. **(D)** Substrates containing a tract of poly(dC)_60,_ or poly(dC)_60_ interrupted every 5^th^ cytosine with either an adenine (C_4_A)_12_ or thymine (C_4_T)_12,_ or poly(dC)_60_ interrupted every 10^th^ cytosine either an adenine (C_9_A)_6_ or thymine (C_9_T)_6_ were replicated. Replication products were analysed on a denaturing agarose gel. **(E)** Replication products from substrates containing a tract of poly(dG)_60_, or (c-MYC Pu22)_8_ or (GGGT)_28_ on either the leading strand template (lead) or the lagging strand template (lag). **(F)** Replication products from substrates containing a tract of poly(dC)_60_, or (DAP)_4_, (DUX4L22)_4,_ or (SNORD112)_4_ on either the leading strand template (lead) or the lagging strand template (lag).

The observed increase in stalling with arrays of quadruplex-forming sequences could either be due to formation of multiple concurrent structures within the same molecule, or due to increased likelihood of structure formation. We noted that poly(dG)_60_, which could in theory form up to four G4 structures, induced more robust stalling than 16 consecutive G4-forming sequences (**Fig. 2A** and **S4A**). In the case of an uninterrupted tract of guanines, a G4 could form from any guanines within the sequence. This is in contrast to other distinct G4-forming sequences we tested (**Fig. 2A**), where structure formation is constrained to a defined window of G-tracts. These observations suggest that stalling efficiency is dictated by the probability of secondary structure formation. To test this possibility, we interrupted the poly(dG)_60_ sequence such that it could still support G4-formation but constrained to specific G-tracts. Leading strand stalling was reduced when the sequence was interrupted, and this was more prominent when interruptions were more frequent (**Fig. 2C**, compare lane 2 to lanes 3 and 4 and **Fig. S4C**). Interrupting the poly(dC)_60_ tract produced similar results (**Fig. 2D** and **Fig. S4D**). We conclude that stalling efficiency is determined by the probability of structure formation.

Having established that replication is stalled due to a quadruplex-forming sequence on the leading strand template, we next sought to determine how a quadruplex-forming sequence on the lagging strand template affects leading strand replication. To this end, we cloned G4-forming sequences such that they were positioned on either the leading or lagging strand template and observed leading strand replication products (**Fig. 2E**). While leading strand stalling was observed when the G4-forming sequence was on the leading strand template, no stalling was seen when it served as the lagging strand template (**Fig. 2E**, compare lanes 3 and 5 to lanes 4 and 6). Although we did not analyse lagging strand synthesis, previous work has demonstrated that the replication machinery skips over a lagging strand G4, resulting in a small gap, about the size of an Okazaki fragment, on the nascent lagging strand (Kumar et al., 2021).

We performed similar experiments to determine how an iM-forming sequence on the lagging strand template affects leading strand synthesis. Interestingly, leading strand stalling occurred in both orientations (**Fig. 2F**). Given the shorter consensus sequence of stable G4s relative to iMs, we speculate that stalling in this scenario is due to G4 formation by the complementary G-rich sequence which is on the leading strand template. In contrast, the C-rich sequences which are complementary to the G4-forming sequences tested are not able to form very stable iM structures (Table 2, both have a low pHT ∼5.8). This may explain why we do not observe leading strand replication stalling when these sequences are positioned on the leading strand template (**Fig. 2E**, lanes 4 and 6).

### CMG can eventually bypass a pre-formed G-quadruplex structure

Having observed consistent replisome stalling at quadruplex-forming sequences, we next wanted to determine whether the stall was transient or persisted over time. To determine this, we carried out pulse-chase experiments, where newly synthesised DNA is labelled with radiolabelled dATP for 10 minutes, after which an excess of unlabelled dATP is added. This prevents labelling of newly initiating replication forks and allows specific analysis of forks that have initiated in the first 10 minutes of the reaction. Replication forks stalled at G4 and iM-forming sequences were not resolved over time and persisted for up to two hours (**Fig. 3A**). This persistent arrest could either be due to blocked unwinding by the CMG helicase or lack of synthesis by polymerase ε. To address the first possibility, we carried out unwinding assays to determine the ability of CMG to unwind pre-formed G4 or iM structures. We chose sequences which induced the strongest replisome arrest – namely (GGGT)_4_ and (SNORD112)_1_, as well as mutated versions which abrogate structure formation (Table 3). These sequences were located on the translocating strand. To favour structure formation, we inserted a poly(dT)_19_ stretch on the opposite strand (Batra et al., 2022). Consistent with previous work (Kumar et al., 2021), time course analysis revealed that CMG unwinding was initially inhibited by a G4 structure, evident within the first 5 to 10 minutes of the reaction (**Fig. 3B**). This was particularly evident when compared to a G4 mutant sequence (**Fig. 3C**), duplex (**Fig. S5A**) or bubble (**Fig. S5B**). However, inhibition of unwinding was not terminal, and CMG was eventually able to unwind G4 substrates to levels similar to the G4 mutated substrate (**Fig. 3B, C**). Interestingly, an iM structure had little effect on CMG unwinding (**Fig. 3D, E**). We observed that unwinding of mutated G4 and iM sequences was slightly less efficient than fully duplexed and bubble substrates (compare **Fig. 3C** and **3E** to **Fig. S5A** and **S5B**). This may be due to interspersed contacts between adenine residues in the mutated sequences and the poly(dT)_19_ loop. To bypass a pre-existing quadruplex structure, CMG may either dismantle the structure or ‘hop’ over and leave it intact. To distinguish between these possibilities, we assessed the effect of a G4-stabilising ligand, PhenDC3 (De Cian et al., 2007). If CMG ‘hops’ over G4s, stabilising them should have no effect. In contrast, if CMG directly unwinds structures, further stabilising a G4 will inhibit unwinding. In the presence of a low concentration of PhenDC3 (0.25 µM), we observed little effect on duplex (**Fig. 3F**) and G4 mutant unwinding (**Fig. S5C**). In contrast, unwinding of the G4 substrate was inhibited (**Fig. 3G**). Therefore, CMG bypasses pre-existing G4 structures by dismantling them rather than ‘hopping’ over them.

**Figure 3.**
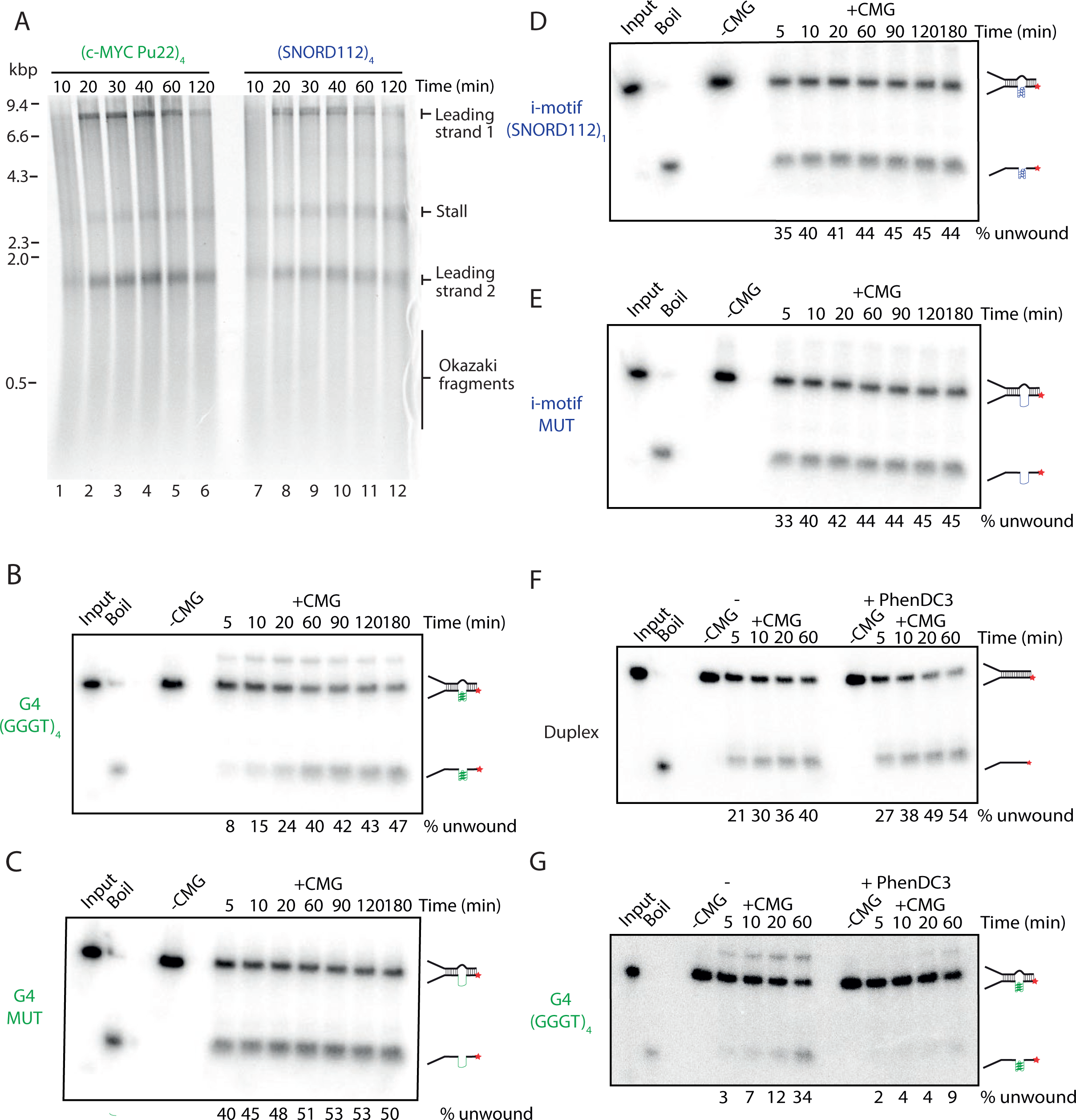
The response of CMG to a pre-formed G4 or iM. (**A**) Pulse-chase time course experiment with (*c-MYC* Pu22)_4_ or (SNORD112)_4_ substrates. Reactions were initiated with radiolabelled dATP for 10 min, chased with excess ‘cold’ dATP and samples taken at the indicated time points. (**B-E**) CMG unwinding assays on substrates containing a pre-formed G4 (B) or a mutant version (see Table 3) (C) or pre-formed iM (D) or a mutant version (see Table 3) (E). CMG unwinding was stimulated by the addition of 2 nM ATP following CMG loading in the presence of ATPγS. Samples were taken at the indicated time points. Products were run on 10% TBE gels. Input and boiled substrates were used as controls to visualise where original and unwound substrates run on the gel. The proportion of template unwound was calculated by measuring the intensity of the ‘unwound’ product band as a proportion of the total product intensity for each lane. (**F-G**) CMG unwinding assays on duplexed substrates (F) or substrates containing a pre-formed G4 (G). Reactions were carried out as in (B-E) but with the addition of 0.25 µM PhenDC3 where indicated.

### Replication templates do not contain pre-formed secondary structures

All the evidence gained thus far indicates that secondary structures are the cause for replication stalling. However, it was not clear whether structures were pre-formed (for example during propagation in bacteria) or were generated during replication. To directly determine whether our replication templates contain pre-existing structures, we employed solid-state nanopore sensing, which was recently utilised to detect quadruplex structures in double-stranded DNA (Boskovic et al., 2019). A positive control contained a single G-quadruplex at a predefined asymmetric position along a dsDNA molecule (Boskovic et al., 2019). DNA is passed through the nanopores and the ionic current measured. Folded G4s give rise to a variation in topology relative to dsDNA that in turn produces a characteristic signal in the ionic current (**Fig. S6A**). The DNA molecule can enter the nanopore in any orientation, which would place the G4 either proximal or distal (**Fig S6B, C**). Besides current drops indicative of G4s, we also observe larger peaks that can be attributed to naturally occurring DNA knots (**Fig. 4A**, ‘knot’). The random position and frequency (∼10%) of these knots is within the expectation for DNA molecules of this length (Plesa et al., 2016).

**Figure 4.**
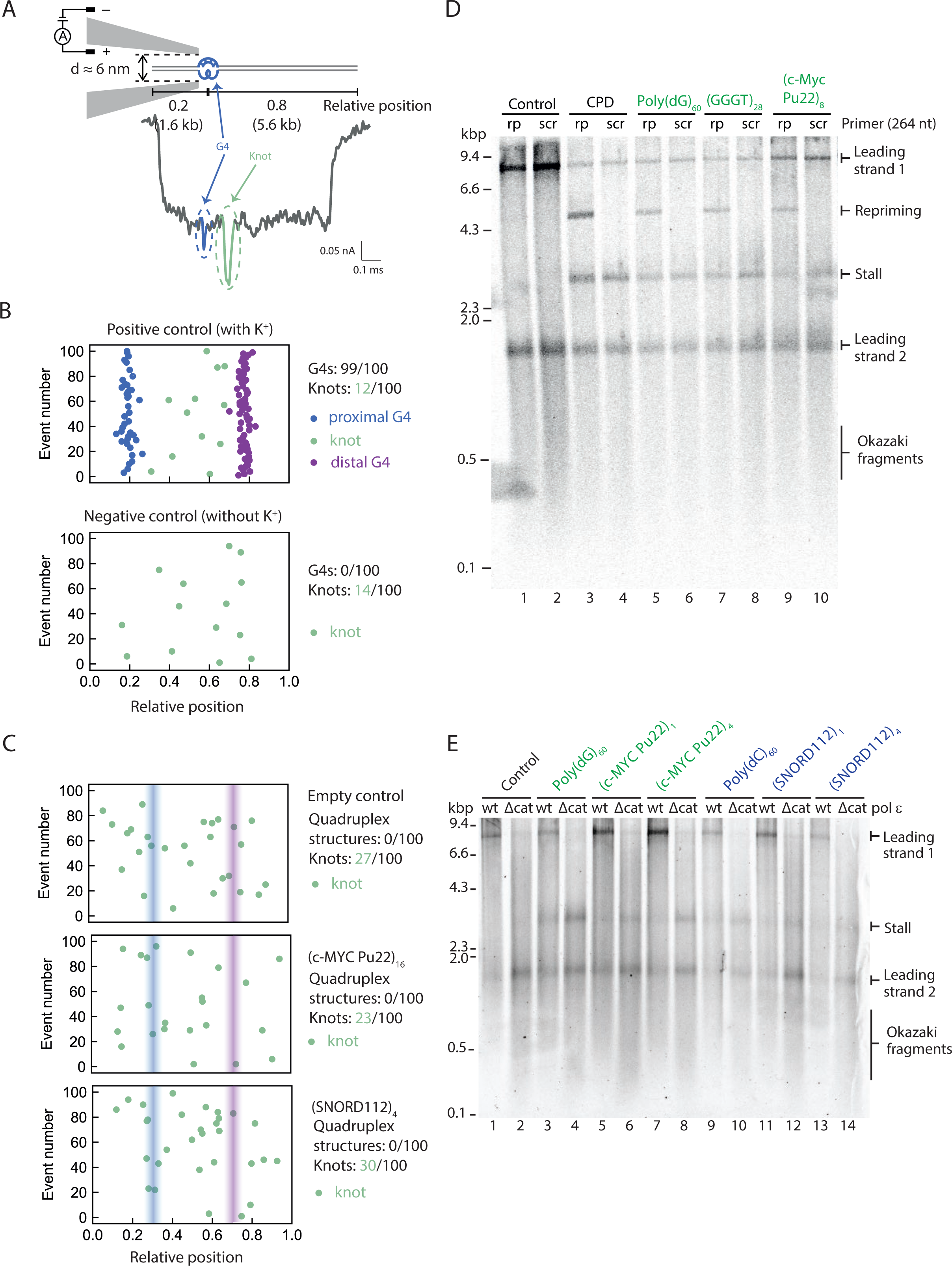
Secondary structures are not pre-formed and likely arise as a result of replication, leading to helicase-polymerase uncoupling. **(A)** Schematic of a positive control DNA passing through the nanopore in the direction that positions the G4 proximally and a representative nanopore measurement event with a DNA knot in the middle of the molecule. The G-quadruplex structure and its corresponding current drop are marked in blue. The DNA knot and its corresponding current drop are marked in green. Numbers indicate the proportion through the DNA the G4 is positioned. **(B)** Summary scatterplots of the peak positions in the first 100 unfolded nanopore events for the positive control (where K^+^ is added) and the negative control (no K^+^ present). The negative control does not contain any G-quadruplex structure without the presence of potassium ions. Numbers indicate proportion of G4s or knots in 100 unfolded events. **(C)** Nanopore measurement results of the first 100 unfolded nanopore events for our replication templates. Summary scatterplots of the peak positions are shown for an empty control, a substrate containing (*c-MYC* Pu22)_16_ and (SNORD112)_4_. Numbers indicate proportion of G4s or knots in 100 unfolded events. **(D)** Replication reactions carried out on G4-containing templates in the presence of a primer that anneals 264 nt downstream of the G4 (rp) or a scrambled control primer (scr). **(E)** Replication of G4 or iM substrates with either *wildtype* pol ε (wt) or a pol ε mutant with a deleted catalytic domain (Δcat). Substrates were digested with EcoRV prior to replication in order to reduce the size of the longer leading strand (leading strand 1). Products were analysed on a denaturing agarose gel.

The summary scatterplot of the peak positions in the first 100 informative events from our nanopore measurement of the positive control is shown in **Figure 4B** (top panel). Virtually all events include a G4 structure (99/100) within the expected relative positions of 0.2 and 0.8. We observed a smaller proportion of naturally occurring knots, randomly distributed along the DNA molecules. Importantly, in the absence of potassium ions, only knots were detected with no discernible G4 signals (**Fig. 4B**, bottom panel).

Having established these positive and negative controls, we next measured our replication substrates, with secondary structures expected at relative positions 0.3 and 0.7 (**Fig. S6D, E**). In contrast to the positive control, none of the replication substrates produced detectable quadruplex peaks. Rather, all substrates, including the empty control, exhibited random knots with a similar distribution and frequency (**Fig. 4C** and **Fig. S6F**). We conclude that our replication substrates do not contain any pre-existing structures.

### Replisome stalling at G4s leads to helicase-polymerase uncoupling

As we had observed that our replication templates do not contain any pre-formed structures and that CMG can directly unwind them, we inferred that they must be forming as a consequence of replication. We hypothesised that structures forming behind CMG inhibit synthesis by polymerase ε, while CMG continues to unwind downstream. This phenomenon, termed helicase-polymerase uncoupling, occurs in response to various leading strand DNA damage lesions (Taylor and Yeeles, 2018) and other types of repetitive sequences (Casas-Delucchi et al., 2022, Kumar et al., 2021) and leads to the exposure of ssDNA. To determine if the same response occurs at G4-forming sequences, we utilised a previously established approach whereby an exogenous primer complementary to the region 264 nt downstream of the G4 sequence is added to the reaction (Taylor and Yeeles, 2018, Casas-Delucchi et al., 2022). This primer can only anneal if ssDNA is exposed, leading to restart of DNA synthesis, thereby serving as a readout for helicase-polymerase uncoupling. As shown in **Figure 4D**, addition of this primer led to the appearance of 5kb restart products for all G4-forming sequences tested, but these were not seen with a scrambled primer. This restart product was absent from the empty vector (**Fig. 4D**, lane 1) but was evident with a template containing a CPD lesion (**Fig. 4D**, lane 3), which served as a positive control. This is strong evidence that unwinding by the CMG helicase is able to continue beyond the G4 sequence, but synthesis by polymerase ε is inhibited. Analysis of the mechanism of stalling at iM sequences using this method is more complex, due to the presence of the intrinsic 5kb band seen in the absence of any exogenous primer upon replication of iM-forming sequences. However, the fact that CMG is easily able to unwind past an iM (**Fig. 3D**), and the evidence that our substrates do not contain any pre-existing structures (**Fig. 4C**), suggests that stalling is due to inhibition of DNA synthesis by pol ε, and as such helicase-polymerase uncoupling may also occur at iM-forming sequences.

Given that our substrates do not contain any pre-formed secondary structures (**Fig. 4C**), we considered the possibility that structures arise on the template ssDNA exposed by CMG unwinding. If this were true, then increasing the amount of ssDNA would enhance the likelihood of structure formation and consequently result in more polymerase stalling. Exposure of excess ssDNA is typically limited as polymerase ε is thought to be tightly coupled to CMG through a direct interaction (Zhou et al., 2017). As pol ε is essential for replication initiation, it cannot be omitted from replication reactions. However, deletion of its catalytic domain, which completely eliminates its polymerase activity, is compatible with replication initiation. Under these conditions, polymerase δ carries out leading strand synthesis. Since polymerase δ does not directly interact with CMG, this results in discontinuous synthesis that is not coupled to unwinding. Replication reactions with this polymerase ε mutant resulted in more replisome stalling when compared to the wildtype protein (**Fig. 4E**, compare odd lanes to even lanes). This was true for both G4 and iM-forming sequences. We conclude that increased replication fork uncoupling leads to more fork stalling and propose this occurs due to increased probability of secondary structure formation.

### Replication products break at i-motifs

Upon replication of substrates containing iM-forming sequences, we consistently observed the presence of a novel 5 kb product on denaturing gels, as highlighted previously. We hypothesised that these products may arise as a result of either re-priming or DNA breakage at the site of the iM during or after replication (**Fig. S1**). To address the latter possibility, we simulated broken replication products by digesting fully replicated control templates post-replication with an enzyme that cleaves within the insert (**Fig. 5A**). The resulting product harbours a double-stranded break at the position of the iM on newly synthesised DNA. To simplify analysis and reduce the heterogeneity in product length arising as a result of flaps generated by strand displacement, pol δ was excluded from these reactions. We verified that this novel 5kb band was unaffected by the presence of pol δ (**Fig. S8C**). Upon analysis by native gel, we observed that the smaller population of products generated by iM substrates migrated at the same positions as the simulated ‘broken products’ (**Fig. 5B**, compare lanes 2 and 3 to lane 4).

**Figure 5.**
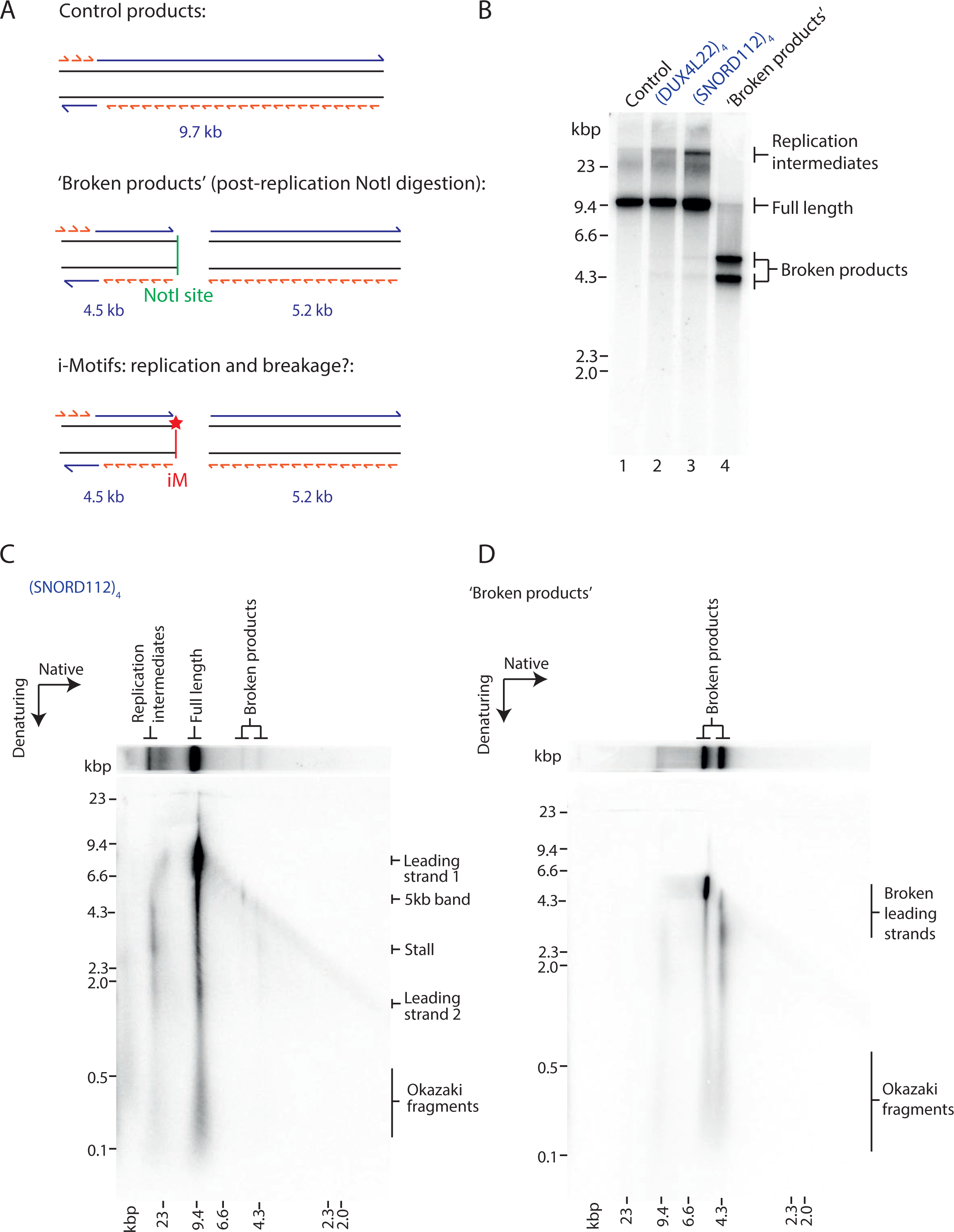
Replication products break at i-motifs. **(A)** Schematic of replication products arising from replication of a control template (top), a control template digested with NotI post-replication to generate ‘broken products’ (middle), and an iM template where potential product breakage occurs at the site of the iM (indicated by a red star) (bottom). **(B)** Visualisation of control or iM-containing replication products on a native gel. ‘Broken products’ was generated by replication of a control template and post-reaction digestion with NotI which cleaves the product at the site of the insert. **(C-D)** Two-dimensional (2D) gel electrophoresis of replication products of an iM-containing template (C) or a ‘broken products’ control (D) (as described in A). Replication products were run firstly in the native dimension and subsequently in the denaturing.

To further analyse these broken replication products, we carried out two-dimensional (2D) gel electrophoresis. As expected, analysis of replication products of an iM template demonstrated the presence of stalled forks and incompletely replicated forks in the population of replication intermediates (**Fig. 5C**). Full-length products consisted mostly of full-length leading strands 1 and 2 and Okazaki fragments. Intriguingly, 2D gel analysis maps the 5 kb band to the products identified on the native dimension as broken (**Fig. 5C**). We also observed a weaker population of products in the denaturing dimension corresponding to the second faster migrating broken product on the native gel (**Fig. 5C**). These smaller products are 1.5 kb and 3-4 kb, which are the expected leading nascent strands within the 4.5 kb reciprocal broken product (**Fig. 5A**). Importantly, these bands mapped to the same positions observed with the simulated broken products (**Fig. 5D**), were absent from the empty vector (**Fig. S7A**) and were also be observed with a different iM-forming sequence (**Fig. S7B**). Together, these results suggest that replication induces breakage within iMs. However, we cannot discount the possibility of intrinsic repriming events (**Fig. S1**), which would migrate as full-length products on the native dimension, but may be masked by the strong signal of leading strand 1 (**Fig. 5C**).

### Stalling at quadruplexes can be rescued by a specialised helicase

Having established the mechanism of fork stalling within quadruplexes, we next wanted to understand how replisomes minimize or resolve stalls. Since increased fork uncoupling enhanced replisome stalling (**Fig. 4E**), we considered the possibility that improved coupling might reduce stalling. CTF18-RFC is an alternative PCNA clamp-loader to the canonical RFC1-RFC that has been proposed to increase coupling of DNA synthesis and unwinding by directly binding polymerase ε (Grabarczyk et al., 2018, Stokes et al., 2020). However, addition of CTF18-RFC, in the absence or presence of RFC1-RFC, did not affect stalling at either G4s or iMs (**Fig. S8A**).

Previous work from our lab had revealed that polymerase δ, as well as high concentrations of dNTPs, can rescue stalling at hairpin-forming sequences, but not at poly(dG)_n_ or poly(dC)_n_ (Casas-Delucchi et al., 2022). Consistent with this, polymerase δ was not able to rescue stalling at any G4 (**Fig. S8B**) or iM-forming sequence (**Fig. S8C**). Similarly, an excess of any dNTP alone, or in combination, was not able to resolve stalling at either G4 (**Fig. S9A**) or iM (**Fig. S9B**) forming sequences. We considered the possibility that the relative ratio of dNTPs may be more important than their absolute concentration. This raised the prediction that an increased proportion of dCTP relative to dA/dG/dTTP might be able to rescue stalling at G4-forming sequences. However, stalled forks were not rescued by a further excess of dCTP (2-fold over dATP and 26-fold over dGTP and dTTP (**Fig. S9C**)). Altogether, this suggests that once the replisome stalls at a quadruplex-forming sequence, the stall is persistent and cannot be overcome by replisome-intrinsic mechanisms.

We previously found that Pif1 could rescue forks stalled at poly(dG)_n_ and poly(dC)_n_ sequences (Casas-Delucchi et al., 2022), raising the possibility that it could rescue stalling at all quadruplex-forming sequences. Pif1 is a well-characterised G4-unwinding helicase shown to play a vital role in enabling the efficient replication of G4s both *in vivo* and *in vitro* (Paeschke et al., 2011, Lopes et al., 2011, Dahan et al., 2018, Sparks et al., 2019b, Byrd et al., 2018, Ribeyre et al., 2009, Paeschke et al., 2013, Maestroni et al., 2020). However, there are additional helicases that bind and unwind G4 structures both *in vitro* and *in vivo*, such as Sgs1 and Chl1. Sgs1 is a RecQ family helicase shown to preferentially unwind G4 DNA and is the yeast homolog of the BLM and WRN helicases (Huber et al., 2002). Chl1 is the yeast homolog of human ChlR1 (also called DDX11). ChlR1 directly unwinds G4 structures and is proposed to help process G4s during DNA replication (Wu et al., 2012, Lerner et al., 2020). To test the role of these helicases, we carried out pulse-chase experiments by pulsing for 10 minutes, during which a persistent stall occurred at G4 or iM-forming sequences. We then added a candidate helicase and allowed replication to continue for a further 10 minutes to determine if the stall could be resolved. Since Chl1 is recruited to replisomes via an interaction with Ctf4 (Samora et al., 2016), we also included Ctf4 in our experiments with Chl1. Neither Sgs1 (**Fig. S10A, S10B**) nor Chl1 (**Fig. S10C, S10D**) were able to rescue stalling at either G4 or iM-forming sequences. Importantly, both of these enzymes were active on a model substrate (**Fig. S10E, S10F**). In contrast, and in agreement with our previous work, Pif1 was able to rescue forks stalled at both G4s and iMs (**Fig. 6A, 6B**). The extent of rescue was less evident for expanded sequences such as (GGGT)_28_ and (SNORD112)_4_ which may indicate that consecutive G4 or iM sequences pose a greater challenge to Pif1. This rescue was dependent on its helicase activity as no rescue was observed with the Pif1 ATPase mutant K264A. The fact that Pif1, but not Sgs1 or Chl1, was able to rescue forks stalled at quadruplex-forming sequences demonstrates the specificity of Pif1, and highlights that only specialised helicases are able to resolve fork stalling at G4 and iM sequences.

**Figure 6.**
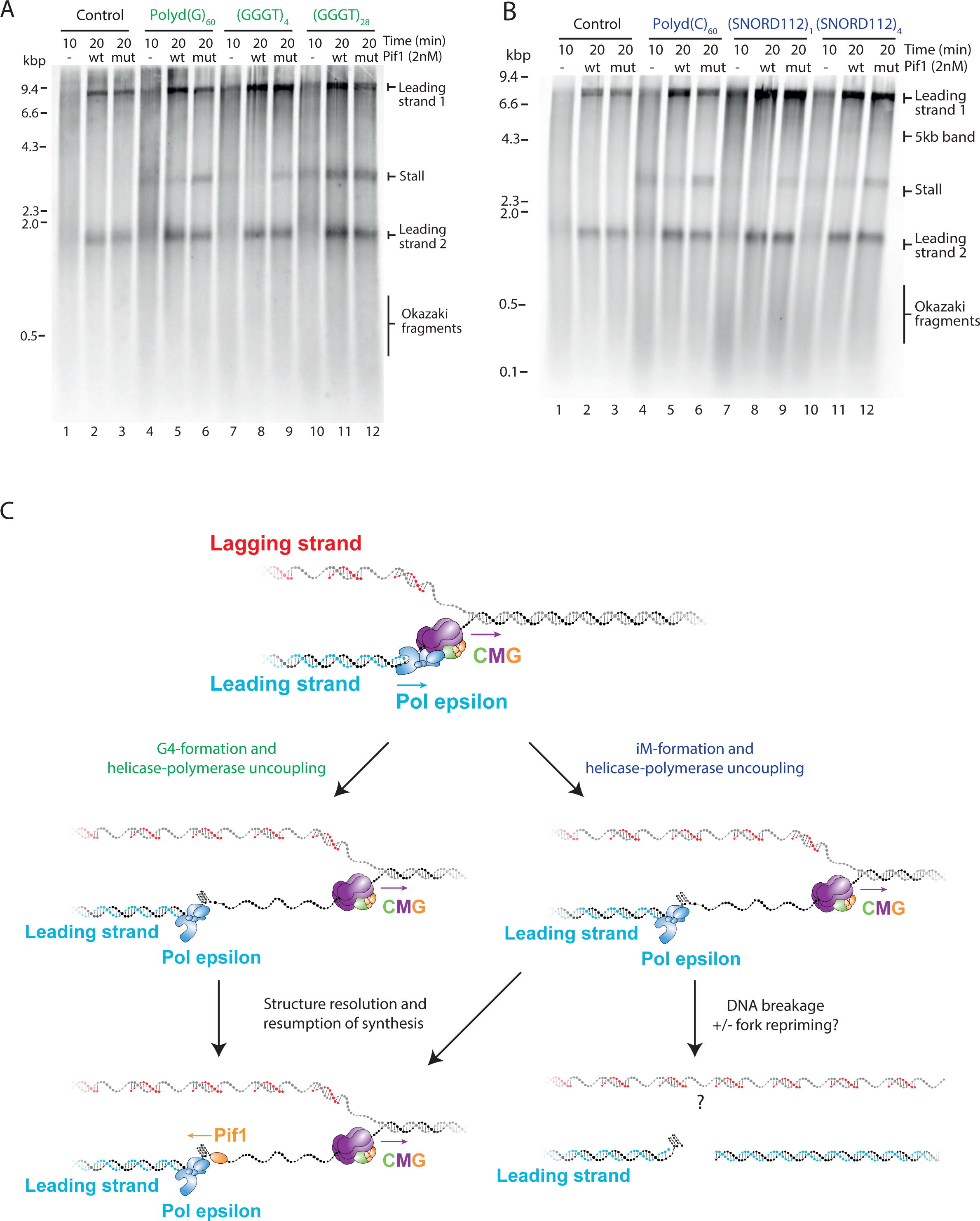
Pif1 is able to rescue stalling at both G4s and i-motifs. (**A** to **B**) Pulse-chase experiments carried out with the indicated templates. Reactions were initiated with radiolabelled dATP. After a 10 min pulse, either *wildtype* or ATPase-dead K264A (mut) pif1 was added with the chase and samples taken after another 10 min. **(C)** Model illustrating the effects of G4s and iMs on replication. CMG unwinds past a leading strand quadruplex-forming sequence and secondary structures form from the exposed ssDNA. These structures inhibit synthesis by polymerase ε, leading to helicase-polymerase uncoupling. Pif1 can unwind both G4s and iMs and allow synthesis to resume. iMs can be resolved in this manner or lead to nascent DNA breakage in the absence or presence of a repriming event. Lagging strand products may remain intact.

## Discussion

We have assessed the response of the eukaryotic replisome to a variety of G4 and iM-forming sequences and found that a single quadruplex-forming sequence alone is able to stall DNA replication. This is a significant finding as these types of sequences are highly prevalent in the human genome. Current estimates are between 370,000 (Huppert and Balasubramanian, 2005, Huppert and Balasubramanian, 2007) and ∼700,000 (Chambers et al., 2015) G4-forming sequences and 5,125 iM-forming sequences (Wright et al., 2017). These must all be replicated accurately in order to maintain genome integrity. Importantly, we have observed that the encounter of a replisome with a single quadruplex forming sequence can lead to the same mechanistic response as triggered in response to DNA damage. Moreover, we have found that in addition to inducing fork stalling, iMs have the propensity to induce DNA breakage. This raises the possibility that there are many physiological DNA sequences within the human genome that have the potential to threaten genome stability.

The ability of a G4-forming sequence to stall DNA replication correlates with its structure-forming potential. Consistent with this, mutated sequences which cannot form structures do not impact replisome progression. Together, these observations provide strong evidence that it is a G4 structure that causes replication stalling, and not the sequence itself. Importantly, we have characterised the effects of iMs on replisome progression. Here, we observed consistent replisome stalling at a variety of physiological iM sequences. Although the response of the replisome to iMs appears largely similar to G4s, we do observe some differences. Firstly, replication stalling at an iM was generally weaker than the stalling we observed at a G4. We speculate that this may be because all of the iMs examined here are weaker structures than those formed by the G4s. This could mean they are less likely to form during replication, although once formed they appear to be a persistent and stable block to replisome progression. Secondly, the ability of iMs to stall replication seems to be less influenced by melting temperature than G4s and may correlate better with transitional pH. Interestingly, previous studies have found that iMs with a higher transitional pH have a higher potential for iM formation *in vivo* and in turn are more likely to undergo mutations and deletions in human cells (Martella et al., 2022). Moreover, the biophysical characterisation of these sequences was done on short oligonucleotides, and the actual thermal stability of these structures within the context of duplexed DNA may in fact be higher. Still, the fact that these structures are more greatly influenced by pH than G4s may explain why we observe a weaker stall in our assays carried out under physiological pH conditions. The fact that stalling occurs in our *in vitro* system under physiological pH strongly supports the idea that iMs are indeed able to form and can be a robust block to the replisome, highlighting their biological relevance.

Importantly, we have observed that iMs can also induce DNA breakage. This is a significant finding as this has the potential to threaten genome stability if not repaired correctly. How and when this DNA breakage occurs in the context of replication remains to be seen. Although we have obtained evidence that a proportion of replication products break at iMs, we cannot rule out the possibility that endogenous re-priming may be occurring preferentially at iM forming sequences. All primases use a purine to initiate primer synthesis, and therefore require a pyrimidine on the template strand. As such, polymerase α has been suggested to prime preferentially at CCC sequences (Davey and Faust, 1990). This may be a mechanism to preserve DNA integrity downstream of forks stalled at iMs, as has been suggested by polymerase α and the 9-1-1 complex downstream of forks stalled at G4s (van Schendel et al., 2021). Another possibility is that the broken products we observe are a consequence of stall-driven repriming events, which somehow promote breakage at iMs. We cannot distinguish whether breakage occurs after unperturbed replication or after stalling and repriming, as both events would yield identical products.

Single molecule solid-state nanopore experiments demonstrate that our substrates do not contain any significant levels of pre-formed structures. Therefore, secondary structures must be forming during replication, resulting in inhibition of synthesis by polymerase ε. As ssDNA is the precursor for structure formation, we favour a model whereby structures form on ssDNA exposed behind the CMG helicase (**Fig. 6C**). This is consistent with a recent super-resolution microscopy study which detected the presence of G4 structures between the CMG helicase and either PCNA or nascent DNA (Lee et al., 2021). Although CMG and polymerase ε are usually tightly coupled in the replisome, it is not altogether clear what the size of the gap is between the exit channel of CMG and the active site of polymerase ε. The most recent structural models of the eukaryotic replisome suggest a gap of at least 16-nt (Yuan et al., 2020). This would be sufficient ssDNA to allow G4 formation, or to nucleate iM formation.

Although we observed consistent replisome stalling at G4 and iM-forming sequences, we never observed a complete block to all replication forks, and only saw a proportion of replication forks stalling. This was true even in the presence of up to sixteen consecutive G4-forming sequences. Therefore, a major determinant of replication fork stalling is the likelihood of structure formation. This explains why we observed only a marginal increase in the proportion of replication forks stalling with an increasing number of consecutive G4s or iMs. In the presence of a larger number of structure-forming repeats, a structure is more likely to fold due to sequence availability, but once a single structure has formed it is sufficient to block the replisome, and additional structures downstream would have no additional effect on synthesis. Similarly, the fact that we consistently observed a greater proportion of replication forks stalling at poly(dG)_60_ and poly(dC)_60_ may be due to the fact that a structure can form in any given window and is not constrained by loop sequences. This is consistent with previous biophysical characterisations of poly(dC)_n_ sequences, which found that the optimum transitional pH peaks at poly(dC)_28_ and gets lower as the number of cytosines increases (up to poly(dC)_40)_) (Fleming et al., 2017). This suggests that iM structures formed by longer tracts of cytosines are not inherently more stable, but rather are more likely to fold. However, once formed, this structure is a robust block to the replisome that cannot be resolved by any replicative polymerase. In addition, structures which fold more quickly may be more likely to fold and in turn stall replication. There are known differences in the kinetics of folding between the two different iM topologies, the 3’E and 5’E conformations, which are distinguished by the position of the outermost C:C^+^ base pair. In the human telomeric iM structure, it has been shown that the 3’E conformation forms faster but this may not be necessarily applicable for all iMs and will depend on the sequences within the loop regions (Malliavin et al., 2003, Lieblein et al., 2013). Folding kinetics usually correlate with structure stability, which may explain why we observe stronger stalling for more thermally stable G4 structures. In addition to the dynamics of the structure itself, the dynamics of how tightly each replication fork is coupled will determine whether there is sufficient time and space for structures to form.

In some cases, G4s and iMs can form on complementary strands of the same sequence. In a physiological context, a stable G4 requires four tracts of three guanines. However, the complementary C-rich sequence would not form a stable iM at a physiological pH. Therefore, a stable G4 does not necessarily equate to a stable iM on the opposite strand. However, a stable iM requires four tracts of five cytosines at a physiological pH (Wright et al., 2017). The complementary G-rich sequence would conform to the requirement of a stable G4, and as such it is more conceivable that a stable G4 structure would be found opposite an iM. This may explain why we observed an orientation-dependent stalling for G4s, but an orientation-independence for iMs.

Similar to our previous work with repetitive sequences (Casas-Delucchi et al., 2022), we observe uncoupling between helicase unwinding and DNA synthesis in response to quadruplexes. This is consistent with previous work which demonstrates an enrichment of G4-forming sequences within 6kb of uncoupled forks in tumour cells (Amparo et al., 2020). This response is akin to the response to leading strand DNA damage (Taylor and Yeeles, 2018) and leads to the exposure of ssDNA. In a physiological context, exposure of large amounts of ssDNA as a result of replication stress can lead to the activation of checkpoint pathways (MacDougall et al., 2007). Further studies are required to explore if such processes occur in response to G4s and iMs during replication. The ssDNA exposed during replisome uncoupling can also be a substrate for recombination and mutation events, which may explain why G4s and iMs are frequent mutation hotspots and undergo rearrangements (Lopes et al., 2011). A recent high-throughput primer extension assay using T7 DNA polymerase demonstrated that polymerase stalling at structure prone sequences generates point mutations with a higher frequency than slippage events. Consistent with their higher propensity to inhibit synthesis, G4s displayed higher mutation rates than iMs and were often mutated in the loop regions (Murat et al., 2020). Whether this same phenomenon occurs with eukaryotic polymerases in the context of the complete replisome remains unclear. Many of these structure-forming sequences have been found at cancer mutation hotspots or breakpoints (Wang and Vasquez, 2017, De and Michor, 2011, Bacolla et al., 2019), and replication stalling could be a potential explanation for mutational events. Indeed, DNA breakage at iMs may directly contribute to their mutagenic potential in cancer.

We have demonstrated that CMG is able to bypass a G4 or iM sequence and established that synthesis by polymerase ε is inhibited. Stabilisation of G4 structures using a small molecule inhibited unwinding by CMG, suggesting that CMG bypasses a G4 by dismantling the structure as opposed to ‘hopping’ over the structure and leaving it intact. This is in contrast with its ability to bypass intact DNA-protein cross-links (Sparks et al., 2019a), leading strand oxidative lesions (Guilliam and Yeeles, 2021) and lagging strand blocks (Langston et al., 2017). It remains to be seen whether replisome stalling occurs in the same position each time. Higher resolution studies could map the position of replisome stalling within the G4 or iM and decipher whether the replisome consistently stalls at the base of the structure, or if it is able to progress some distance through.

It is unsurprising that we observe rescue of replication fork stalling at G4s by Pif1, given the breadth of data describing its role as a G4-unwinding helicase (Paeschke et al., 2013, Paeschke et al., 2011, Lopes et al., 2011, Dahan et al., 2018, Sparks et al., 2019b, Byrd et al., 2018, Ribeyre et al., 2009, Maestroni et al., 2020). However, the fact that Pif1 is also able to resolve forks stalled at iMs points towards the broad specificity of Pif1 as a helicase. This is in line with its ability to also resolve forks stalled at hairpins (Casas-Delucchi et al., 2022). Our observation that rescue of replisome stalling was less efficient with an array of consecutive G4 and iM sequences may indicate that multiple structures are forming within these arrays which may require more extensive helicase activities.

Interestingly, none of the other helicases tested were able to rescue fork stalling at G4s or iMs, despite their demonstrated ability to unwind G4s (Huber et al., 2002, Wu et al., 2012). This may reflect the fact that these helicases show a preference for certain types of G4s. For example, Chl1 has been shown to have stronger unwinding activity on anti-parallel G4s (Wu et al., 2012), while the G4 sequences we tested all form parallel G4s which is a more common conformation. This may also explain the large number of quadruplex-unwinding helicases that appear to have some level of redundancy, as they may each have a role in resolving structures in different scenarios. For example, BLM helicase has been shown to suppress recombination at G4s in transcribed genes (van Wietmarschen et al., 2018). The fact that only an accessory helicase was able to resolve stalls at G4s and iMs may also explain why mutations in helicases such as these lead to genome instability diseases, such as Bloom’s Syndrome (reviewed in (Cunniff et al., 2017)), Werner’s syndrome (Yu et al., 1996) and Fanconi Anaemia (reviewed in (Brosh and Cantor, 2014)).

Recent advances in the field have provided the tools to reconstitute human DNA replication *in vitro* (Baris et al., 2022). Although the core replication machinery is conserved from budding yeast to human, it remains to be seen whether G4s and iMs have the same impact on progression of the human replisome. Using this system to study replication of structure-forming sequences would also enable one to study the roles of other human proteins in this process, such as an array of human helicases including RecQ helicases such as BLM (Sun et al., 1998) and WRN (Fry and Loeb, 1999), and the Fanconi Anaemia protein FANCJ (Wu et al., 2008).

Altogether, we discovered that a range of physiological G4 and i-motif structures arising as a result of DNA replication can stall the eukaryotic replisome. This study provides further insight as to why these sequences provide a barrier to DNA replication and suggests a potential mechanism for structure formation and resolution during replication. Moreover, we have found that i-motifs can directly cause DNA breakage. We therefore propose that endogenous DNA secondary structures are a source of replication stress, which may explain their genomic instability and mutation frequencies in cancer.

## Materials and Methods

### Constructing repeat-containing replication substrates

Substrates for replication assays were constructed by inserting repetitive sequences into a 9.8kb parental plasmid (pGC542) containing a single synthetic yeast replication origin (ARS306) which has been previously described (Casas-Delucchi et al., 2022). Repetitive sequences were cloned into the MCS 3kb downstream of the origin and expanded using a strategy that employs synthetic oligos and type IIS restriction enzymes (Scior et al., 2011). All oligos were ordered from Integrated DNA Technologies (IDT), the sequences of which can be found in Table 3.

### Preparing templates for replication assays

Plasmids (Table 4) were transformed into NEB Stable Competent *E. coli* cells (#C3040I) which are ideal for propagation of repeat-containing plasmids. Cultures were grown at 30°C to reduce potential recombination and mutation events. Plasmids were purified using a QIAGEN HiSpeed Maxi Kit. Subsequently, supercoiled plasmids were isolated from nicked plasmids by PlasmidSelect clean up. DNA samples were diluted 6-fold in 100 mM Tris-HCl (pH 7.5), 10 mM EDTA and 3 M (NH_4_)_2_SO_4_ (final concentration = 2.5 M) before incubation with 300 µL PlasmidSelect Xtra slurry (pre-washed with 100 mM Tris-HCl (pH 7.5), 10 mM EDTA and 2.3 M (NH_4_)_2_SO_4_) for 30 minutes to bind. Following binding, nicked plasmids were eluted initially with 1 ml buffer consisting of 100 mM Tris-HCl (pH 7.5), 10 mM EDTA and 1.9 M (NH_4_)_2_SO_4_. This was repeated twice. Then, 1 ml buffer was added to the beads and allowed to incubate for 10 minutes at RT. Subsequently, supercoiled plasmids were eluted with 100 mM Tris-HCl (pH 7.5), 10 mM EDTA and 1.5 M (NH_4_)_2_SO_4_ by incubation for 10 minutes at RT. This was repeated once. Samples were de-salted by dialysis against 0.1xTE for 3 hours and overnight. DNA was concentrated using a 100 kDa Amicon concentrator followed by ethanol precipitation and resuspended in 1xTE.

### CPD substrate

The substrate containing a site-specific DNA damage (CPD lesion) was prepared as has been previously described (Casas-Delucchi et al., 2022).

### Protein purification

All protein expression strains, and expression and purification steps were carried out as previously described (Casas-Delucchi et al., 2022).

#### CMG

*Saccharomyces cerevisiae* CMG was expressed from the budding yeast strain yJCZ3 (Zhou et al., 2017). All purification steps were carried out as previously described (Baretic et al., 2020), with the following exceptions. After elution from Calmodulin Sepharose 4B (GE healthcare), the eluate was applied to a MonoQ 5/50 GL column (GE Healthcare) equilibrated in 25 mM Tris-HCl pH 7.2, 10% glycerol, 0.005% TWEEN 20, 0.5 mM DTT, 150 mM KCl. The protein was eluted with a 20 CV gradient from 150-1000 mM KCl and peak fractions were dialysed overnight against 500 ml 25 mM HEPES-KOH pH 7.6, 40 mM KOAc, 40 mM K-glutamate, 2 mM Mg(OAc)2, 0.25 mM EDTA, 0.5 mM DTT, 40% glycerol. The protein was subsequently concentrated through a 0.5 ml 30K MWCO Pierce™ Protein Concentrator (Thermo Scientific, 88502), snap-frozen and stored at −80°C.

#### Pol ε-Δcat

The DNA polymerase ε mutant with a deleted catalytic domain (Pol ε-Δcat) was expressed from the budding yeast strain yAJ25, which has been previously described (Yeeles et al., 2017), and purified as per wildtype pol ε.

#### Sgs1

Purified recombinant yeast Sgs1 was expressed and purified as previously described (Cejka and Kowalczykowski, 2010).

#### Chl1

Chl1 expression strain

Y5562 MAT**a**, ade2-1, ura3-1, his3-11,15, trp1-1, leu2-3,112, can1-100, pep4::HIS3, ade2::ADE2-pRSII402-Gal4-pGal1/10-Chl1-FLAG-Protein A

Chl1 purification

Budding yeast cells overexpressing Chl1 were grown in YP medium containing 2% raffinose as the carbon source to an optical density of 1.0 at 30°C. 2% galactose was then added to the culture to induce the protein expression, and the cells were further grown for 90 minutes. Cells were collected by centrifugation, washed with deionized water and suspended in Chl1 buffer (50 mM Tris-HCl pH 7, 10% glycerol, 2 mM MgCl_2_, 0.5 mM TCEP) containing 0.1% Triton X-100, 500mM NaCl, 0.5mM Pefabloc, as well as the cOmplete-EDTA protease inhibitor cocktails. The cell suspension was frozen in liquid nitrogen, then cells were broken in a cryogenic freezer mill. The cell powder was thawed on ice, and further Chl1 buffer containing 0.1% Triton X-100 and 500 mM NaCl, and protease inhibitors was added. The lysate was clarified by centrifugation at 20,000 x g for 1 hour. The clarified lysate was transferred to pre-equilibrated IgG agarose beads. 8 μg/ml RNase A (Merck) was added, and incubated for 2 hours. The resin was washed with Chl1 buffer containing 0.1% Triton X-100 and 500 mM NaCl and then incubated in Chl1 buffer containing 0.1% Triton X-100, 500 mM NaCl, 10 mM MgCl2 and 1 mM ATP for 15 minutes. The resin was washed again with Chl1 buffer containing 0.1% Triton X-100 and 500 mM NaCl and incubated overnight in the same buffer containing 10 μg/ml PreScission protease. The eluate was collected, and Chl1 dilution buffer (50 mM Tris-HCl pH7, 10% glycerol, 2 mM MgCl_2_, 0.5 mM TCEP, 10 mM NaCl) was added to adjust the salt concentration to 160 mM NaCl. The diluted sample was loaded onto a HiTrap Heparin (Cytiva) column, equilibrated with Chl1 buffer containing 160 mM NaCl. The column was developed with a linear gradient from 160 mM to 1 M NaCl in Chl1 buffer. The peak fractions were pooled and loaded onto a Superdex 200 Increase (Cytiva) gel filtration column that was equilibrated and developed with Chl1 gel filtration buffer (20 mM Tris-HCl pH 7.5, 150 mM NaCl, 10% Glycerol, 0.5 mM TCEP). The peak fractions were concentrated by ultrafiltration.

#### RFC-Ctf18

RFC-CTF18 expression strain: yEF4

MATa ade2-1 ura3-1 his3-11,15 trp1-1 leu2-3,112 can1-100

bar1::Hyg

pep4::KanMX ura3::URA3pRS306/Rfc2, CBP-Rfc3

trp1::TRP1pRS304/Rfc4, Rfc5

his3::HIS3pRS303/Ctf18

leu2::LEU2pRS305/Ctf8, Dcc1

Expression and purification steps were carried out as per RFC purification (Casas-Delucchi et al., 2022).

### In vitro replication assays

MCM loading was carried out for 10 minutes at 24°C on 3nM circular DNA that was linearised during loading with 0.3 µL AhdI in a buffer containing 25 mM HEPES (pH 7.6), 10 mM magnesium acetate, 100 mM potassium glutamate, 1 mM DTT, 0.01% NP-40-S, 0.1 mg/ml BSA, 80 mM potassium chloride, 5 mM ATP, 20 nM ORC, 45 nM Cdc6, 75 nM Cdt1-Mcm2-7 and 50 nM DDK. Loading was stopped by the addition of 120 nM S-CDK for 5 minutes at 24°C. Following loading, samples were diluted in a buffer containing 25 mM HEPES (pH 7.6), 10 mM magnesium acetate, 100 mM potassium glutamate, 1 mM DTT, 0.01% NP-40-S, 0.1 mg/ml BSA to dilute the final contribution of chloride to 14 mM. A nucleotide mix was added to give final concentrations of 200 μM ATP, CTP, GTP and UTP; 30 μM dATP, dCTP, dGTP and dTTP; and 132 nM α-P^33^-dATP. Subsequently, to initiate replication, a master mix of proteins was added to give final concentrations of 100 nM GINS, 10 nM S-CDK, 10 nM Mcm10, 40 nM Csm3/Tof1, 20 nM Pol ε, 30 nM Dpb11, 40 nM Cdc45, 40 nM Mrc1, 60 nM RPA, 40 nM RFC, 120 nM PCNA, 5 nM Pol δ, 50 nM Pol α, 20 nM Sld3/7 and 20 nM Sld2. Reactions were incubated at 30°C for 40 minutes. For samples to be run on denaturing gels, 0.5 µl SmaI was added to each 10 µl reaction in the final 10 minutes of the reaction, which cleaves products approx. 100 bp from the origin of replication. This removes heterogeneity in the length of leading strand products arising due to variability in the exact location where synthesis of leading strands begins, despite origin specificity (Taylor and Yeeles, 2018). Reactions were quenched by adding EDTA to a final concentration of 100 mM.

Pulse-chase experiments were carried out as previously described (Casas-Delucchi et al., 2022). During the pulse, unlabelled deoxyribonucleotide concentrations were as follows: 30 μM dCTP, dTTP, dGTP and 2.5 μM dATP. To carry out the chase, unlabelled dATP was added to a final concentration of 400 µM, with the addition of 400 µM of dGTP, dCTP or dTTP, or 800 µM dCTP where indicated.

Repriming experiments were carried out by the addition of 60 nM oligonucleotide after loading and before initiation of replication (Casas-Delucchi et al., 2022).

### Post-replication sample processing

For denaturing gels, after quenching with 20mM EDTA, 1/10 volumes of alkaline loading dye (0.5M NaOH, 10% sucrose, xylene cyanol in water) was added to samples. Replication products were separated on 0.8% alkaline agarose gels in 30 mM NaOH, 2 mM EDTA at 32V for 16 hours. Subsequently, products were fixed on denaturing agarose gels by incubation in 5% TCA for 40 minutes at room temperature.

For native gels, replication products were treated with 0.1% SDS and 1/100 volumes of Proteinase K at 37°C for 20 minutes. DNA was extracted using phenol:chloroform:isoamyl alcohol 25:24:1 (Sigma-Aldrich, P2069) and samples were subsequently passed over illustra MicroSpin G-50 columns (Sigma-Alrich, GE27-5330-02) to remove incorporated nucleotides and exchange the buffer to TE. 6× NEB™ Purple Gel Loading Dye (no SDS) (NEB, B7025S) was added to samples before separation on a 0.8% agarose/TAE gel in 1xTAE at 30V for 22 hours at 4°C.

For two-dimensional (2D) gel electrophoresis, samples were treated as per native gels and then split equally into two lanes on the same native gel as described above (one lane for analysis and one lane for the second dimension). One lane was excised from the native gel and soaked for 2 × 1 hour in alkaline running buffer. Gel slices were then horizontally inserted into the top of a 0.8% alkaline agarose gel and run as per standard denaturing gels.

Gels were dried before being exposed to a Storage Phosphor Screen (GE Healthcare, BAS-IP MS 2025) and imaged on a Typhoon scanner (Cytiva). Images were analysed and quantified in ImageJ. Quantifications of stalling intensities were calculated from an average of three replicates in separate experiments. To calculate intensities, the background for each lane was subtracted from each measurement. The 3kb stalling intensity band was normalised to the intensity of ‘leading strand 2’ in each lane to account for variation in the efficiencies of reactions for each substrate.

### Preparing templates for helicase assays

Forked DNA substrates were prepared as previously described (Batra et al., 2022). Briefly, dried oligos from IDT were resuspended to 10 µM in 10 mM Tris pH 8.0. The bottom strand of the substrate (GC340, SW051 or SW105) was end labelled with γ-^32^P-ATP in a reaction containing 5 pmol oligo, 1X PNK buffer, 1U of PNK enzyme (NEB, M0201S), and γ-^32^P-ATP (0.03 mCi). The reaction was incubated for 1 hour at 37°C, followed by heat inactivation of PNK for 20 minutes at 80°C and then 10 minutes at 90°C. Unincorporated radiolabelled nucleotides were removed by passing the sample through a G50 column (GE healthcare, 2753002) equilibrated in 10mM Tris pH 8.0. The concentration of K+ was then adjusted to 50 mM by the addition of 1 M KOAc. To anneal the complementary top strand (GC339 or SW114), 1 µl of a 10 µM stock of oligonucleotide was added and the reaction was incubated at 95°C for 5 minutes, followed by a slow cooling to 10°C at a rate of −1°C/minute. Annealed products were run on 10% TBE gels (ThermoFisher Scientific, EC62755BOX) in 0.5xTBE at 150 V for 45 minutes. Fully annealed substrates were isolated from the gel using a crush-soak method as described in Batra et al., 2022.

For Chl1 and Sgs1 unwinding assays, substrates containing a 5’ overhang (for Chl1) and a 3’ overhang (for Sgs1) were generated as previously described (Casas-Delucchi et al., 2022).

Sequences of all oligonucleotides used to generate helicase unwinding substrates are detailed in Table 3.

### Helicase assays

CMG unwinding assays were carried out using 0.5 nM of labelled substrate and 20 nM of purified CMG. Reactions were assembled in a buffer containing 25 mM HEPES pH 7.6, 10 mM MgOAc, 30 mM NaCl, 0.1 mg/ml BSA and 0.1 mM AMP PNP. Reactions were incubated at 30°C for 30 minutes to allow CMG to pre-load onto the template, before addition of ATP to a final concentration of 2 mM to stimulate unwinding. At this point, 65 nM of the unlabelled version of the labelled oligo (GC340, SW051, SW069, SW105, SW129 or SW130) was added to trap any unwound oligos and prevent substrate re-annealing. Reactions were incubated at 30°C and time points taken as indicated. Reactions were stopped by the addition of 0.5% SDS and 200 mM EDTA, supplemented with Novex Hi-Density TBE Sample buffer (ThermoFisher Scientific, LC6678) and analysed on 10% Novex TBE gels (ThermoFisher Scientific, EC62755BOX) in 1× TBE at 90 V for 90 minutes. Gels were exposed to a Storage Phosphor Screen (GE Healthcare, BAS-IP MS 2025) and imaged in a Typhoon (Cytiva). Images were analysed and quantified in ImageJ.

For Chl1 and Sgs1 unwinding assays, reactions were carried out using 0.5 nM of labelled duplex and 50 nM of purified recombinant protein as previously described (Casas-Delucchi et al., 2022).

### Biophysical characterisation of G4 sequences

#### DNA Annealing Step

DNA sequences (Table 1) were purchased in their lyophilised form with standard desalting purification from Integrated DNA Technology (IDT) and re-dissolved in MilliQ water to reach a stock concentration of 100 µM. The sequences were then further diluted to 10 µM in the annealing buffer (25 mM HEPES, 10 mM MgCl_2_, 110 mM KCl/LiCl), heated to 95℃ for 15 minutes and left to slowly cool down to room temperature overnight.

#### Thermal difference spectra (TDS)

100 µL of each of the 10 µM DNA sequences (annealed as above) was transferred into a High Precision Cell (quartz glass) Light Path 10 mm (HellmaAnalytics) and covered with 200 µL of mineral oil to prevent evaporation. The cuvettes were sealed with a plastic lid and transferred into the Agilent Cary 3500 UV-Vis Multicell Peltier spectrometer. A first scan was run at 25℃ (Scan range = 800-200 nm | Averaging time (s) = 0.02 | Data Interval (nm) = 1 | Scan rate (nm/min) = 3000 | Spectral bandwidth (nm) = 2). The samples were then heated to 95℃ and left to equilibrate for 7 mins before running a second scan. Each TDS curve was obtained by subtracting the absorbance spectra (25°C) by the absorbance spectra (95 °C).

#### UV-vis melting curve

100 µL of each of the 10 µM DNA sequences (annealed as above) was transferred into a High Precision Cell (quartz glass) Light Path 10 mm (HellmaAnalytics) and covered with 200 µL of mineral oil to prevent evaporation. The cuvettes were sealed with a plastic lid and transferred into the Agilent Cary 3500 UV-Vis Multicell Peltier spectrometer. The melting curve was obtained by imputing the following parameters (Wavelengths: 295 nm | Averaging time (s) = 0.1 | Spectral bandwidth (nm) = 2) and the heating protocol in Table 5. Data was collected only for stages 5 and 6 for each run.

**Table 5.** Heating protocol for Uv-Vis melting curve of DNA sequences.

| Stage | Data Interval (°C) | Rate (°C/min) | Temperature Start/End (°C) |
| --- | --- | --- | --- |
| 1 | 1 | 5 | 25/95 |
| 2 | 1 | 5 | 95/25 |
| 3 | 1 | 5 | 25/95 |
| 4 | 1 | 5 | 95/25 |
| 5 | 0.1 | 0.5 | 25/95 |
| 6 | 0.1 | 0.5 | 95/25 |

T_m_ was extrapolated for each sample by fitting the relative melting curve as shown by Mergny and Lacroix (Mergny and Lacroix, 2003) with the Python script provided by Giacomo Fabrini (Fabrini, 2022).

### Biophysical characterisation of i-motif sequences

#### Oligonucleotides

DNA sequences (Table 2) were supplied by Eurogentec (Belgium), synthesized on a 1000 nmol scale and purified by reverse phase HPLC. All DNA sequences were dissolved in ultra-pure water to give 100 μM final concentrations, which were confirmed using a Nanodrop. For all experiments, ODNs were diluted in buffer containing 10 mM sodium cacodylate and 100 mM potassium chloride at the indicated pH. DNA samples were thermally annealed by heating in a heat block at 95°C for 5 minutes and cooled slowly to room temperature overnight.

#### Circular Dichroism

CD spectra were recorded on a Jasco J-1500 spectropolarimeter using a 1 mm path length quartz cuvette. ODNs were diluted to 10 µM (total volume: 100 μL) in buffer at pH increments of 0.25 or 0.5 pH units from 4.0 to 8.0, depending on the sequence. Spectra were recorded at 20 °C between 200 and 320 nm. Data pitch was set to 0.5 nm and measurements were taken at a scanning speed of 200 nm/min, response time of 1 s, bandwidth of 2 nm and 100 mdeg sensitivity; each spectrum was the average of four scans. Samples containing only buffer were also scanned according to these parameters to allow for blank subtraction. Transitional pH (pH_T_) for each iM was calculated from the inflection point of fitted ellipticity at 288 nm.

#### UV Absorption Spectroscopy

UV spectroscopy experiments were performed on a Jasco J-750 equipped with a Julabo F-250Temperature Controller and recorded using low volume masked quartz cuvettes (1 cm path length). Annealed DNA samples (250 μL) were transferred to a cuvette and covered with a stopper to reduce evaporation of the sample. The absorbance of the DNA was measured at 295 nm and 260 nm as the temperature of the sample was held for 10 min at 4°C, heated to 95°C at a rate of 0.5°C per min, then held at 95°C for 10 min before the process was reversed; each melting/annealing process was repeated three times. Data were recorded every 1°C during both melting and annealing and melting temperatures (T_m_) were determined using the first derivative method. TDS were obtained by subtracting the spectrum of the folded structure between 220 and 320 nm at 4°C from that of the unfolded structure at 95°C. The data was normalized and maximum change in absorption was set to +1 as previously described (Mergny et al., 2005).

#### Data Analysis

Final analysis and presentation of the data was performed using GraphPad Prism version 9.0. All sets of data passed Shapiro-Wilk normality test, p-values were calculated by One way ANOVA followed by Holm-Sidak posthoc analysis for the melting temperature and thermodynamic data collected from the triplicate measurements for each oligonucleotide.

### Nanopore detection of structures

Plasmids for nanopore analysis were pre-linearised at the origin of replication by digestion with SmaI. This generated linear templates with the structure-forming sequence positioned asymmetrically from the ends of the DNA (30% into the template, 70% from the other end of the template). Following digestion, DNA was extracted using phenol:chloroform:isoamyl alcohol 25:24:1 (Sigma-Aldrich, P2069) and subsequently ethanol precipitated and resuspended in ddH_2_O.

#### Synthesis of control DNA molecules

The linear ssDNA scaffold for our designs as the positive control and negative control was obtained by cutting circular M13mp18 DNA (New England Biolabs) at BamHI-HF and EcoRI-HF (New England Biolabs, 100 units/μL) restriction sites following our published protocol (Bell and Keyser, 2016). Staple 42 and staple 43 in the basic staple set (Bell and Keyser, 2016) were replaced by our customized strand G16 with four repeats of GGGT in the middle to form the G-quadruplex secondary structure in the presence of potassium cations. Another strand, cG16, was added to form a double helix with the scaffold at the position where the G-quadruplex formed and thus stabilized the structure. All the DNA oligonucleotides were purchased from Integrated DNA Technologies, Inc. (IDT). Detailed sequences of the customized oligonucleotides can be found in Table 6. After mixing the modified oligonucleotide set and the M13mp18 scaffold at a 5:1 stoichiometric ratio, the solution was heated to 70°C followed by a linear cooling ramp to 25°C over 50 minutes. Excess oligonucleotides were removed using Amicon Ultra 100kDa filters. After quantification with Nanodrop 2000 spectrophotometer, samples were then kept in a freezer at −20°C for later measurements.

**Table 6.**
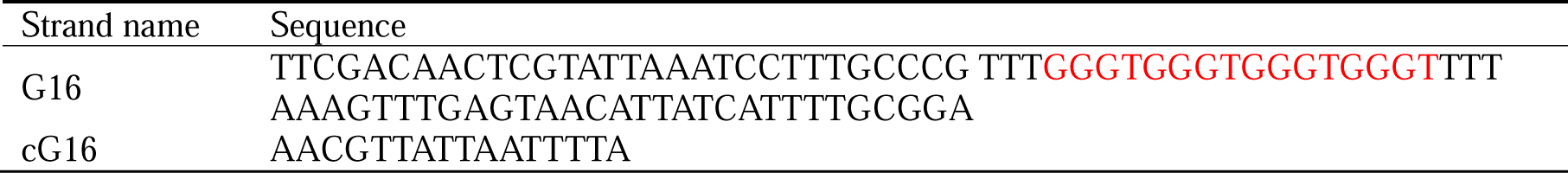
Detailed sequences of the customized oligonucleotides.

#### Nanopore measurement

Nanopores used in this project were fabricated by laser-assisted pulling (P-2000, Sutter Instrument) of quartz capillaries (outer diameter 0.5 mm and inner diameter 0.2 mm, Sutter Instrument) as in our previous work (Bell and Keyser, 2016), though a higher pulling temperature (HEAT=500) were used to obtain smaller nanopores with diameters of about 6 nm and higher signal-to-noise ratio. Current-voltage characteristic curves from −600 mV to 600 mV were recorded to indicate the estimated sizes and the root-mean-square (RMS) noise of the ionic current through the nanopores. Details can be found in Table 7. Once functional nanopores had been identified, the central reservoir of our nanopore chip was filled with our DNA sample (0.2 nM in 200 mM KCl and 4 M LiCl buffer, except for 0.2 nM in 4 M LiCl buffer when measuring the negative control group). A positive voltage of 600 mV was then applied to drive negatively charged DNA molecules through the nanopore, creating characteristically transient changes in the ionic current trace. Data analysis was performed with the LabVIEW software and self-written python programs. It is important to note that a small slope was often observed at the baseline, probably due to the slight change in the buffer concentration as the measurement was carried out. The current baseline was linearly fitted, and the slope was corrected in Figure 4A-4C and Figure S6 for better presentation. When we drew a reference line at 0.15 nA below the current baseline, the two intersections of the reference line and the current trace of one event were taken as its start and end. The event duration Δ is the timescale between the start and end, and Δ refers to the interval between any second-level peak beyond the first-level plateau and the start (Figure S6A). The position of the peak is calculated as 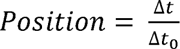.

**Table 7.** Parameters of the nanopores used.

| DNA sample | Diameter (nm) | Root-mean-square noise (pA) |
| --- | --- | --- |
| Positive control | 6.3 | 6.0 |
| Negative control | 5.9 | 6.3 |
| Empty control | 6.3 | 5.8 |
| 16×Myc | 6.0 | 5.4 |
| 55×GGGT | 6.1 | 5.7 |
| 60×G/C | 5.7 | 6.0 |
| 4×SONRD | 6.1 | 5.7 |
| 4×DUX4L22 | 5.3 | 5.6 |

## Author Contributions Statement

S.L.W. and C.S.C.D. performed all experiments, except for biophysical characterisations of structures and nanopore experiments. G.C. conceptualised and supervised the research and acquired funding. S.L.W. wrote the manuscript with input from G.C., M.D.A., F.R., Z.A.E.W., D.G., Y.L. and U.F.K. M.D.A. and F.R. have designed and performed biophysical characterisation of G4-structures. Z.A.E.W. and D.G. have designed and performed biophysical characterisation of i-motif-structures. U.F.K. and Y.L. performed solid-state nanopore experiments to detect quadruplex structures. M. M. expressed and purified recombinant Chl1. E. E. F. and J. T. P. Y. generated the expression strain for recombinant RFC-CTF18.

## Acknowledgements

This work was funded by a Wellcome Trust and Royal Society Sir Henry Dale fellowship (210470/Z/18/Z) as well as internal funding from the Institute of Cancer Research. Marco Di Antonio is supported by a Biotechnology and Biological Sciences Research Council (BBSRC) David Phillips Fellowship [BB/R011605/1] and a Lister Prize Fellowship. Federica Raguseo is supported by a Leverhulme Trust, Cellular Bionics scholarship [EP/S023518/1]. Dilek Guneri is supported by a BBSRC grant [BB/W001616/1]. We would like to thank Petr Cejka (Institute for Research in Biomedicine, Switzerland) for providing published purified recombinant protein. We would like to thank Max Douglas for reagents and experimental support, and Manuel Daza Martin for critical reading of the manuscript. We are thankful to Giacomo Fabrini and Lorenzo Di Michele for support on Tm analysis. U.F.K and Y.L. thank F. Boskovic for help with the design of the G4 control structures and help with sample preparation for the nanopore experiments.

