## Supplementary Figures for "Replication-induced DNA secondary structures drive fork uncoupling and breakage"

### Figure S1

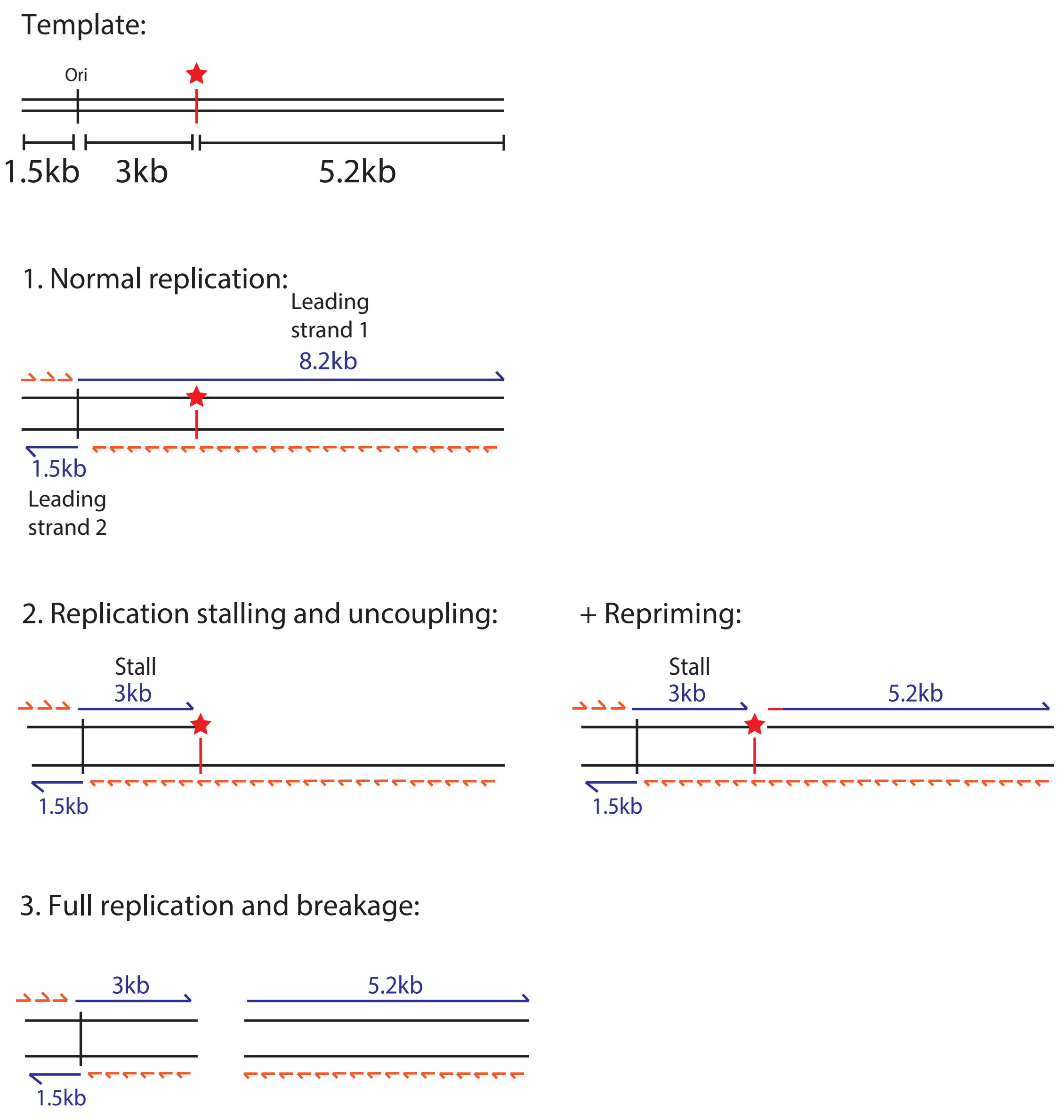


**Figure S1. Schematic of potential replication products from G4 and iM-containing templates.** Position of the origin of replication is marked as 'ori' from which replication initiates through sequence-specific replisome loading. Two forks initiate from the origin and generate one longer rightward moving fork of 8.2kb ('Leading strand 1') and one shorter leftward moving fork of 1.5kb ('Leading strand 2'). Lagging strand products are synthesised as short Okazaki fragments. The multiple cloning site (indicated by a red star) is positioned 3kb downstream of the origin and was used to insert various G4 or iM-forming sequences into the template. This means that only the rightward moving fork would encounter them and therefore 'leading strand 2' can serve as an internal control. Under conditions of replication stalling, helicase-polymerase uncoupling may occur which may be associated with intrinsic repriming at the site of fork stalling. Replication products may also break at the site of the insert during or after replication.

Figure S2

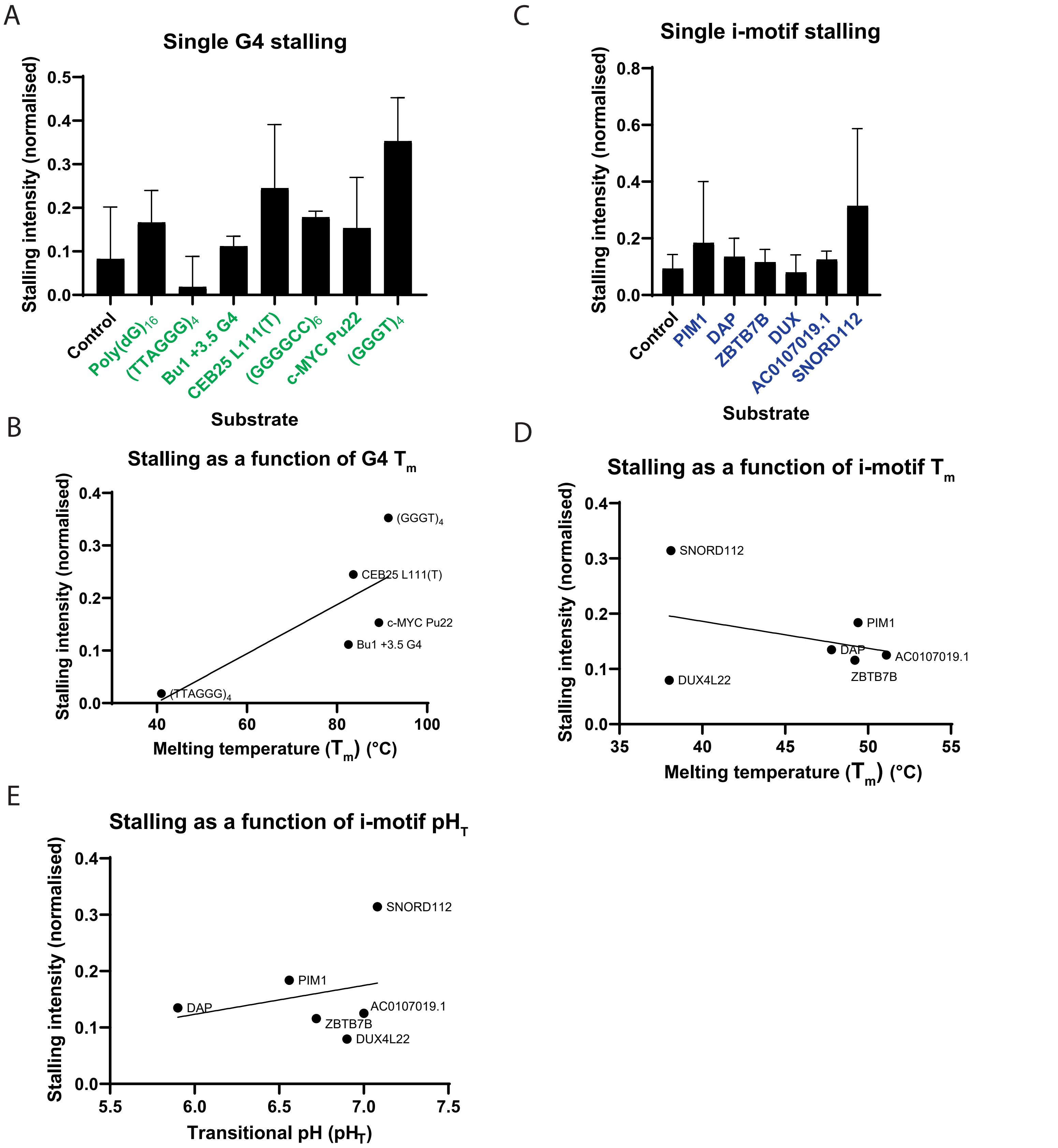


**Figure S2. Quantification of replisome stalling at G4s and iMs in relation to their biophysical characteristics.** (A-B) Quantification of stall intensities from three independent experiments of substrates, as shown in Fig. 1B and 1C. The 3kb stalling intensity band was normalised to the intensity of 'leading strand 2' in each lane to account for variation in the efficiencies of reactions for each substrate. The mean stalling intensity is plotted with error bars representing the standard error. (C) Melting temperatures of the G4s formed by sequences tested in Fig. 1B versus their stalling intensities as calculated in (A). Line indicates simple linear regression with a Pearson correlation r value of 0.76. (D) Melting temperatures of the iMs formed by sequences tested in Fig. 1C versus their stalling intensities as calculated in Fig. S2B. Line indicates simple linear regression with Pearson correlation r value of -0.34. (E) The transitional pH (pHT) of the iMs formed by the sequences tested in Fig. 1C versus their stalling intensities as presented in (B). Line indicates simple linear regression with a Pearson correlation r value of 0.27.

### Figure S3

A

| G4 mutated substrates | Sequence (5' - 3') |
|---|---|
| *c-MYC* Pu 22 | TGAGGGTGGGTAGGGTGGGTAA |
| *c-MYC* Pu 22 MUT | TGAGAGTGAGTAGAGTGAGTAA |
| $(GGGT)_4$ | GGGTGGGTGGGTGGGT |
| $(GGGT)_3$ | GGGTGGGTGGGT |
| CEB25 L111(T) (Piazza et al., 2015) | AAGGGTGGGTGGGTGGGTGTGAGT GTGGGTGTGGAGGTAGATGT |
| CEB25 L111(A) (Piazza et al., 2015) | AAGGGAGGGAGGGAGGGTGTGAG TGTGGGTGTGGAGGTAGATGT |

B

| iMotif mutated substrates | Sequence (5'-3') |
|---|---|
| DUX4L22 | CCCCCGAAACGCGCCCCCCTCCCCCCTC CCCCCTCTCCCCC |
| DUX4L22 MUT | CCTCCGAAACGCGCCTTCCTCCTTCCT CCTTCCTCTCCTCC |
| SNORD112 | CCCCCCCCCGCCCCCCACCCCCCCACCCC CCCCCCC |
| SNORD112 MUT | CCTCCTTCCGCCTTCCACCTTCTCACC TCCTCCTCC |

E

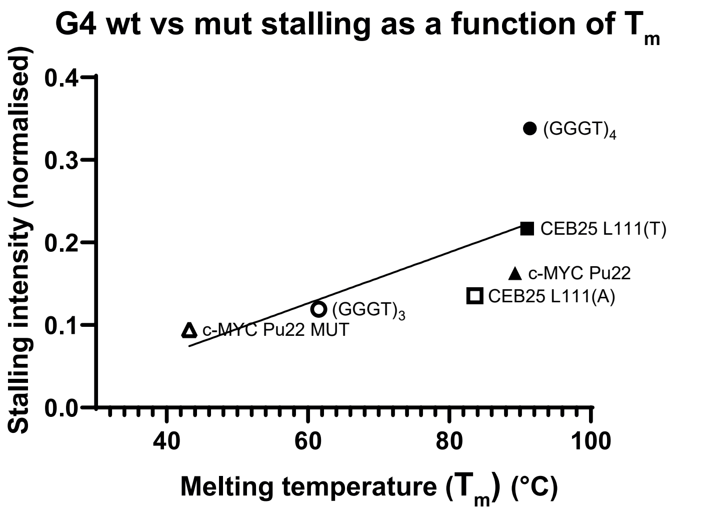


F

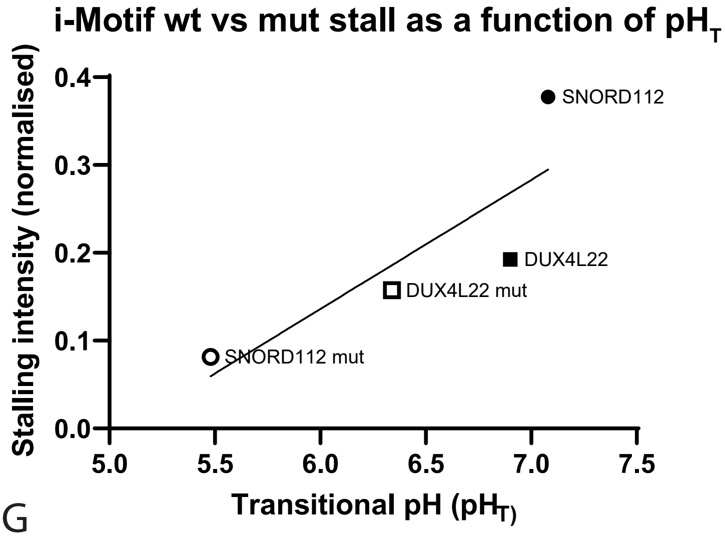


C D

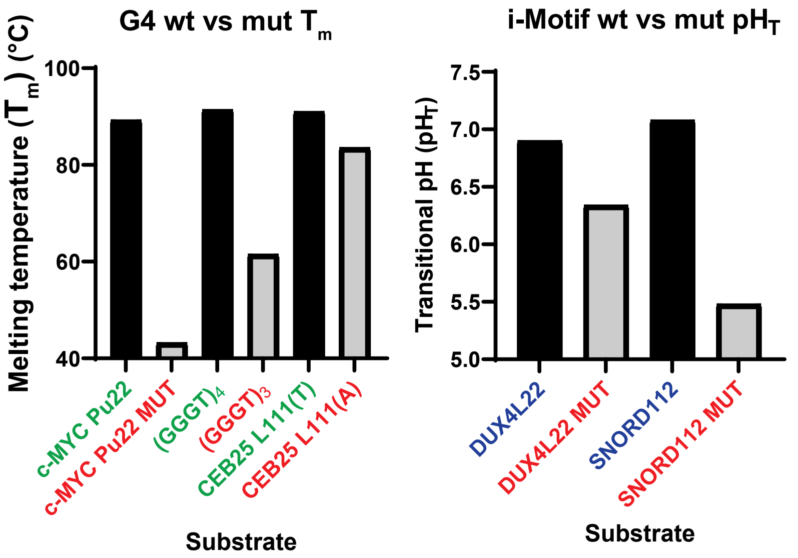


G

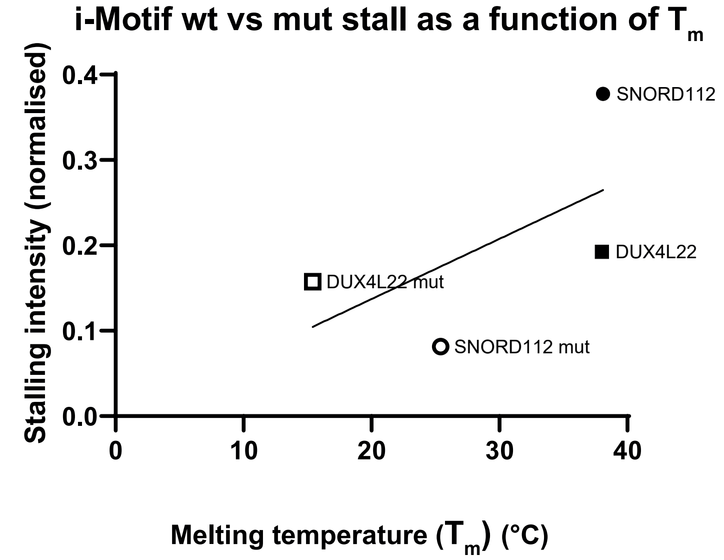


**Figure S3. Correlations between biophysical properties of mutated G4 and iMs and replisome stalling.** (A and B) Sequences of wildtype or mutated G4 (A) or iM-forming (B) sequences used to generate substrates for replication reactions in Fig. 1D and 1F. Mutated sequences (depicted in red) have abrogated or weakened ability to form structures. (C) Melting temperatures of the secondary structures formed by sequences depicted in Fig. 1D. (D) The transitional pH (pHT) of the structures formed by the sequences depicted in Fig. 1F. (E) Melting temperatures of the structures formed by sequences tested in Fig. 1D versus their stalling intensities as calculated in Fig. 1E. Line indicates simple linear regression with Pearson correlation r value of 0.7. (F and G) Transitional pH ($pH_T$) (F) or melting temperature (G) of the structures formed by sequences tested in Fig. 1F versus their stalling intensities as calculated in Fig. 1G. Lines indicate simple linear regression with Pearson correlation r values of 0.8 (F) and 0.6 (G).

Figure S4

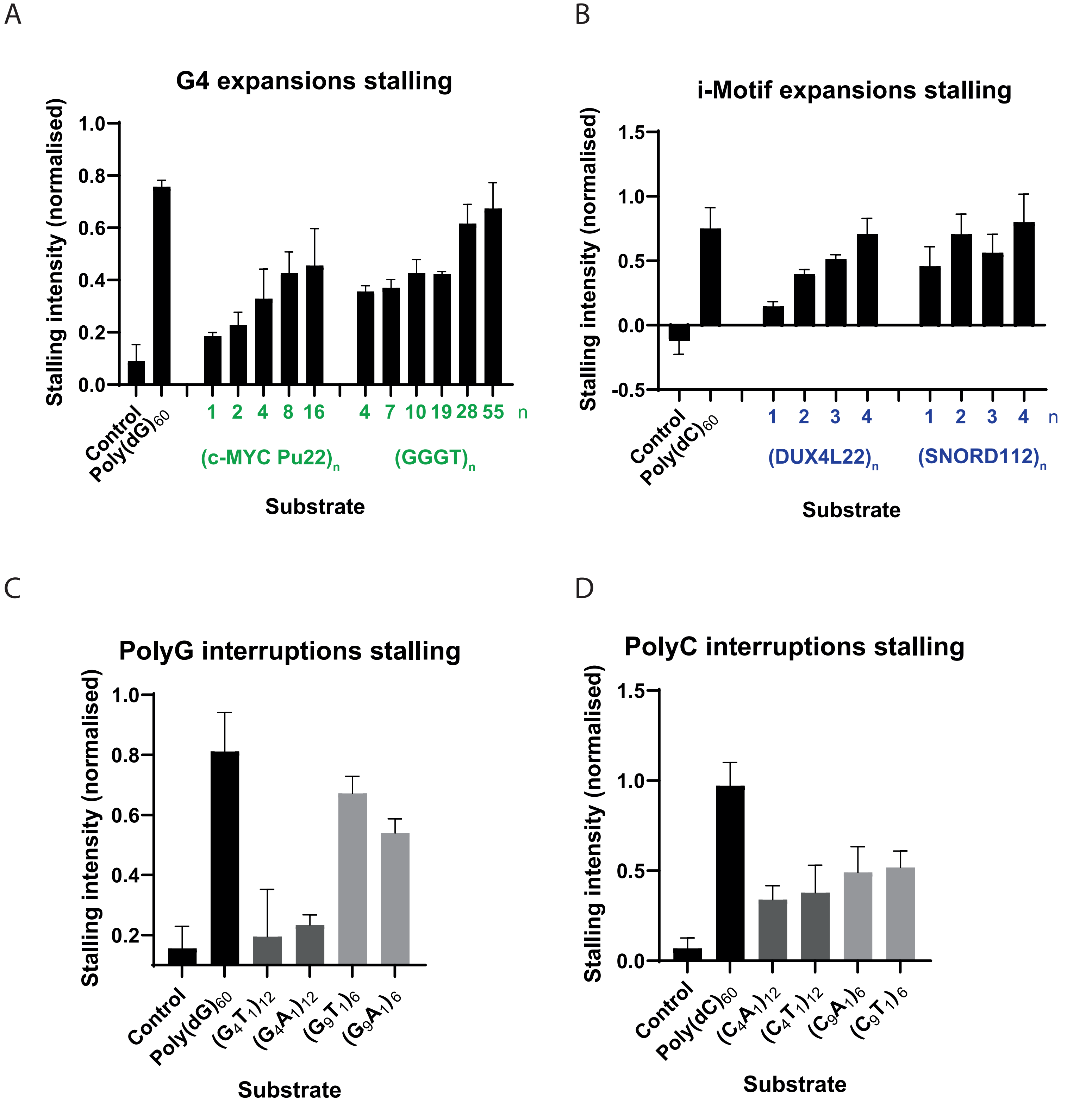


**Figure S4. The effect of increased consecutive quadruplex sequences and interrupting tracts of poly(dG)$_{60}$ or poly(dC)$_{60}$ on replisome stalling.**
(A-D) Quantification of stall intensities from three independent experiments of relevant substrates, as indicated in Fig. 2A-D. The 3kb stalling intensity band was normalised to the intensity of ‘leading strand 2’ in each lane to account for variaton in the efficiencies of reactions for each substrate. The mean stalling intensity is plotted with error bars representing the standard error.

Figure S5

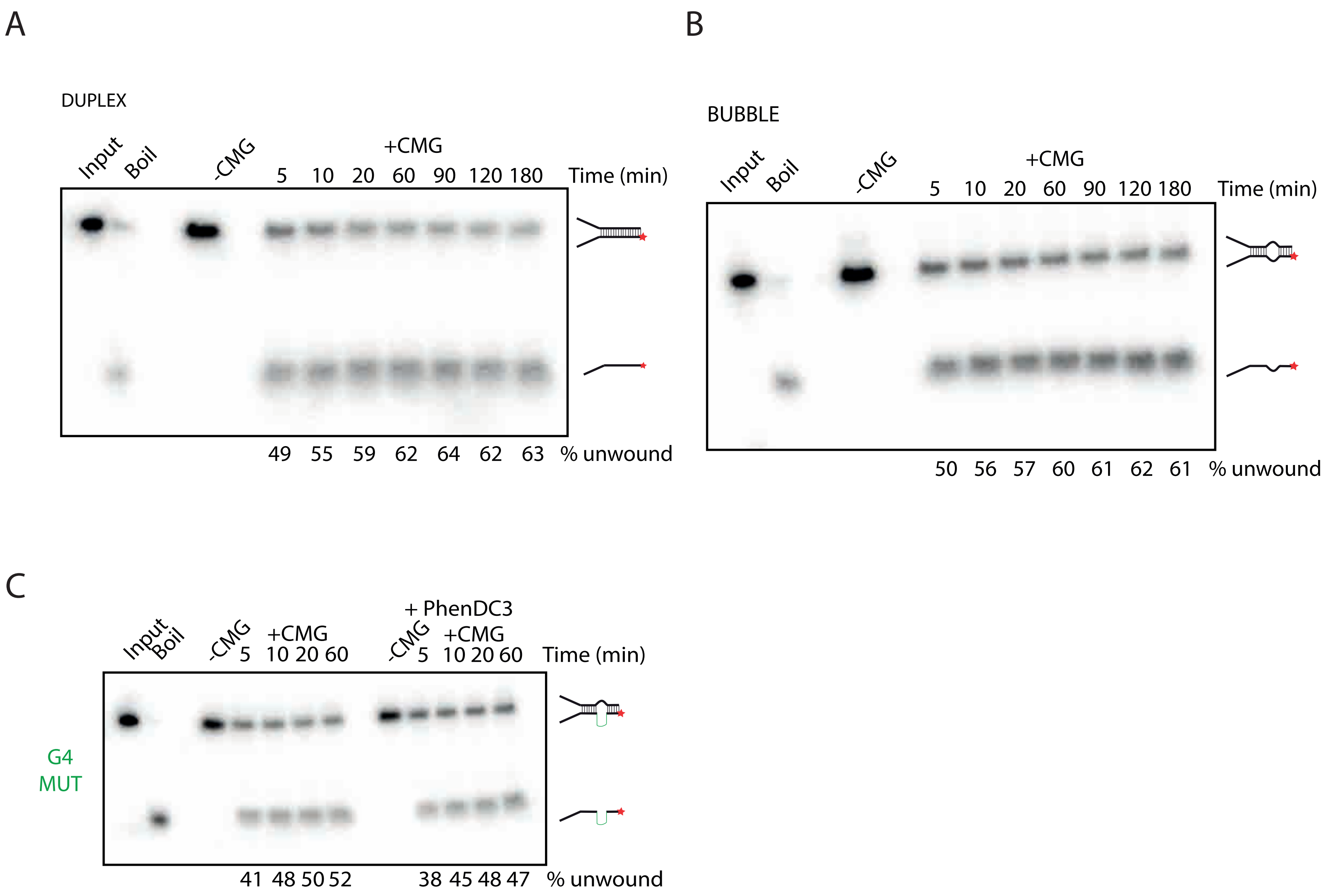


**Figure S5. Unwinding of non-structure containing substrates by CMG.**
(A-B) CMG unwinding assays on substrates containing duplex (A) or a poly(dT)$_{19}$ bubble (B). CMG unwinding was stimulated by the addition of 2 nM ATP following CMG loading in the presence of ATPγS. Samples were taken at the indicated time points. Products were run on 10% TBE gels. Input and boiled substrates were used as controls to visualise where original and unwound substrates run on the gel. The proportion of template unwound was calculated by measuring the intensity of the ‘unwound’ product band as a proportion of the total product intensity for each lane. (C) CMG unwinding assays on substrates containing G4 mutant sequence. Reactions were carried out as in (B) but with the addition of 0.25 μM PhenDC3 where indicated.

#### Figure S6

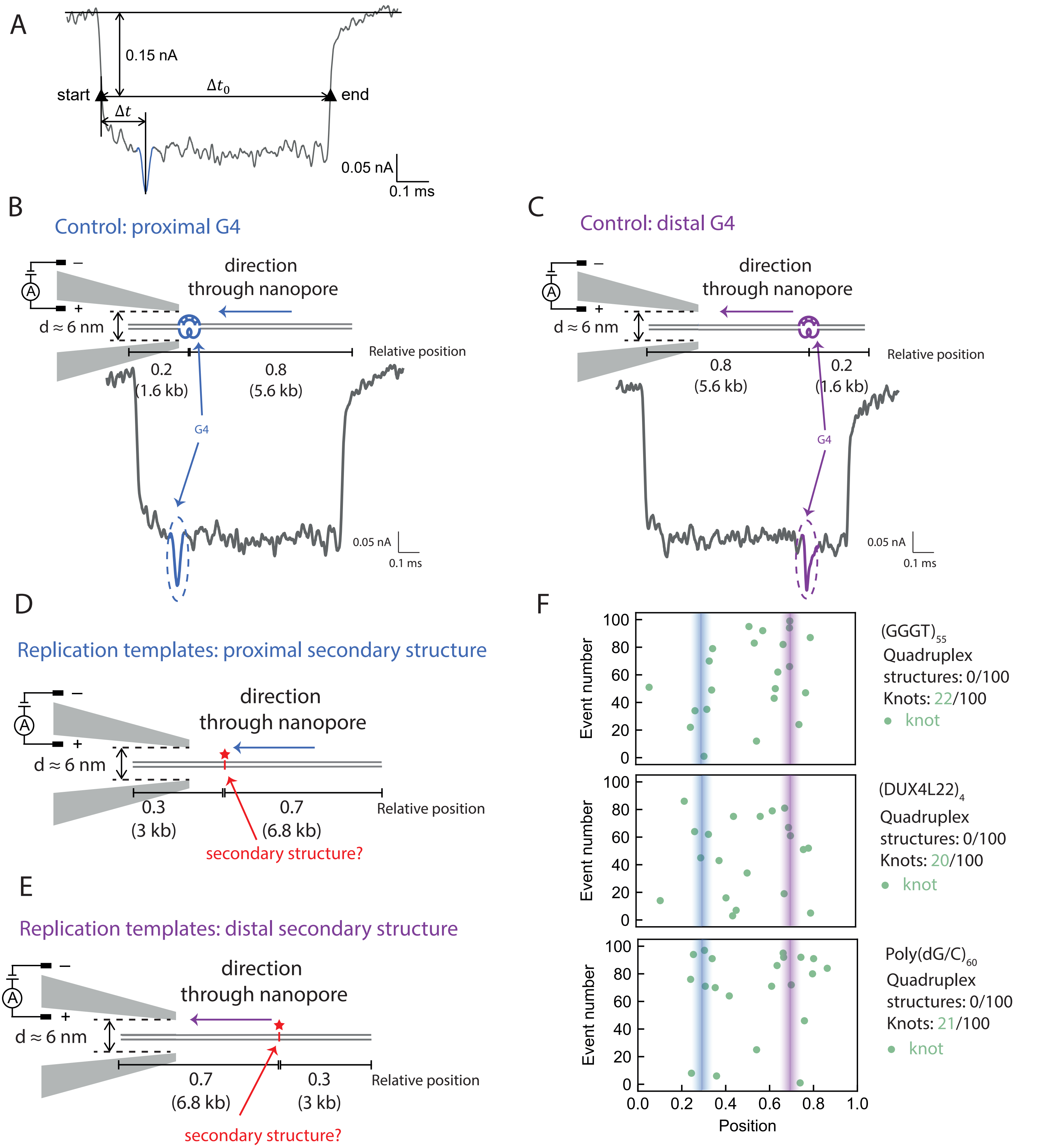


**Figure S6. Single molecule nanopore experiments to detect secondary structures.**
(A) A representative event of our nanopore measurements. The two black triangles, namely the two i ntersections of the reference line 0.15 nA below the current baseline and the event current trace, mark the start and end of one event. 〖Δt〗_0 refers to the timescale between the start and end, and Δt refers to the interval between second-level peak and the start. (B and C) Schematic of a positive control DNA passing through the nanopore in the direction that positions the G4 proximally (B) or distally (C), and a representative nanopore measurement event. Numbers indicate the proportion through the template the G4 is positioned. The G-quadruplex structure and its corresponding current drop are marked in blue (B) or purple (C).
(D and E) Schematic of DNA replication templates passing through the nanopore in the direction that positions the potential secondary structures proximally (D) or distally (E). Numbers indicate the proportion through the template the secondary structure-forming sequence is positioned. (F) Nanopore measurement results of $(GGGT)_{55}$, $(DUX4L22)_4$ and Poly$(dG/C)_{60}$. Lines indcate expected positions of secondary structures.

### Figure S7

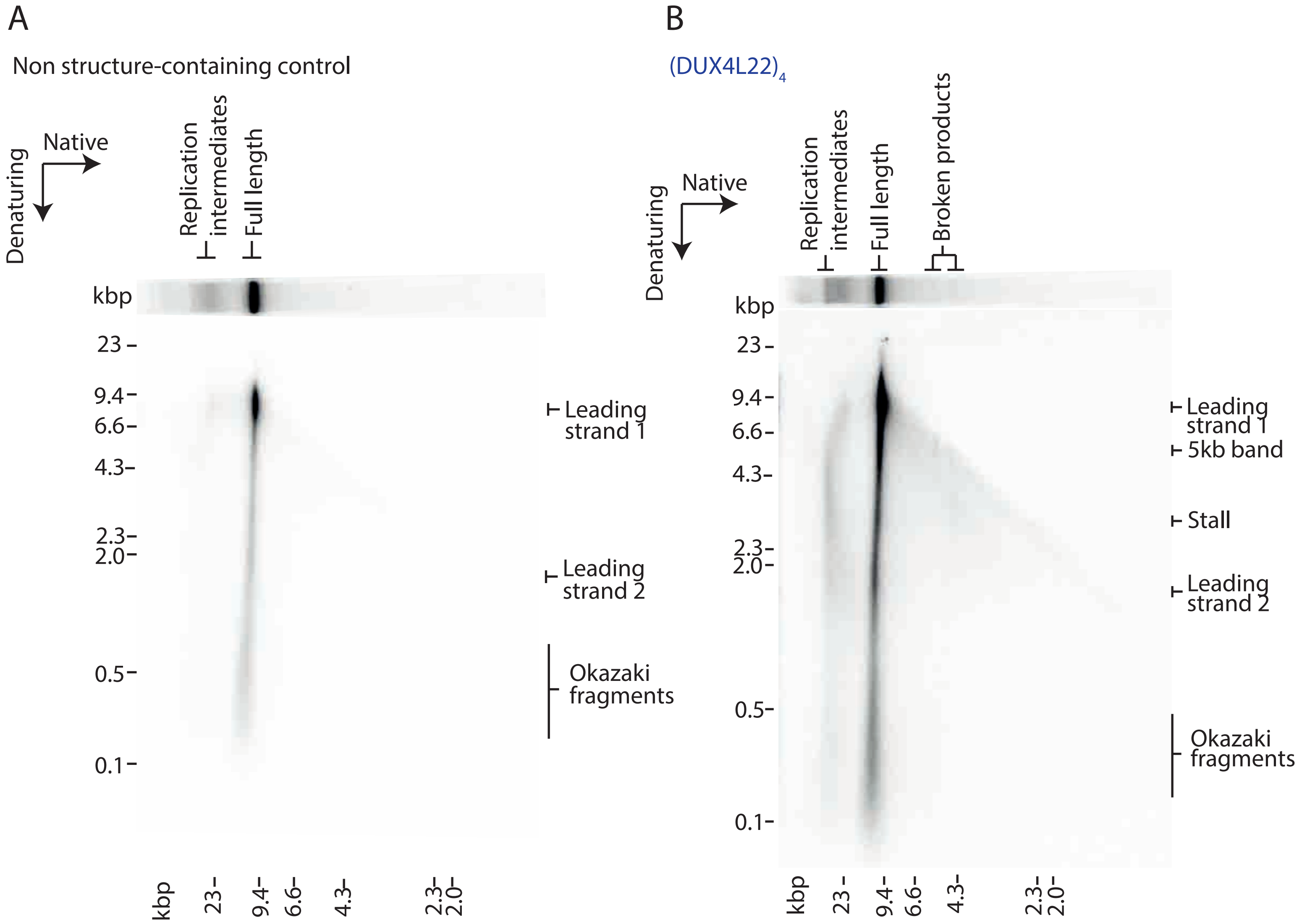


**Figure S7. Analysis of replication products by two-dimensional (2D) gel electrophoresis.**
(A and B) Analysis of replication products of control (A) or iM (B) substrates by 2D gels. Replicated products were run firstly in the native dimension and subsequently in the denaturing dimension.

Figure S8

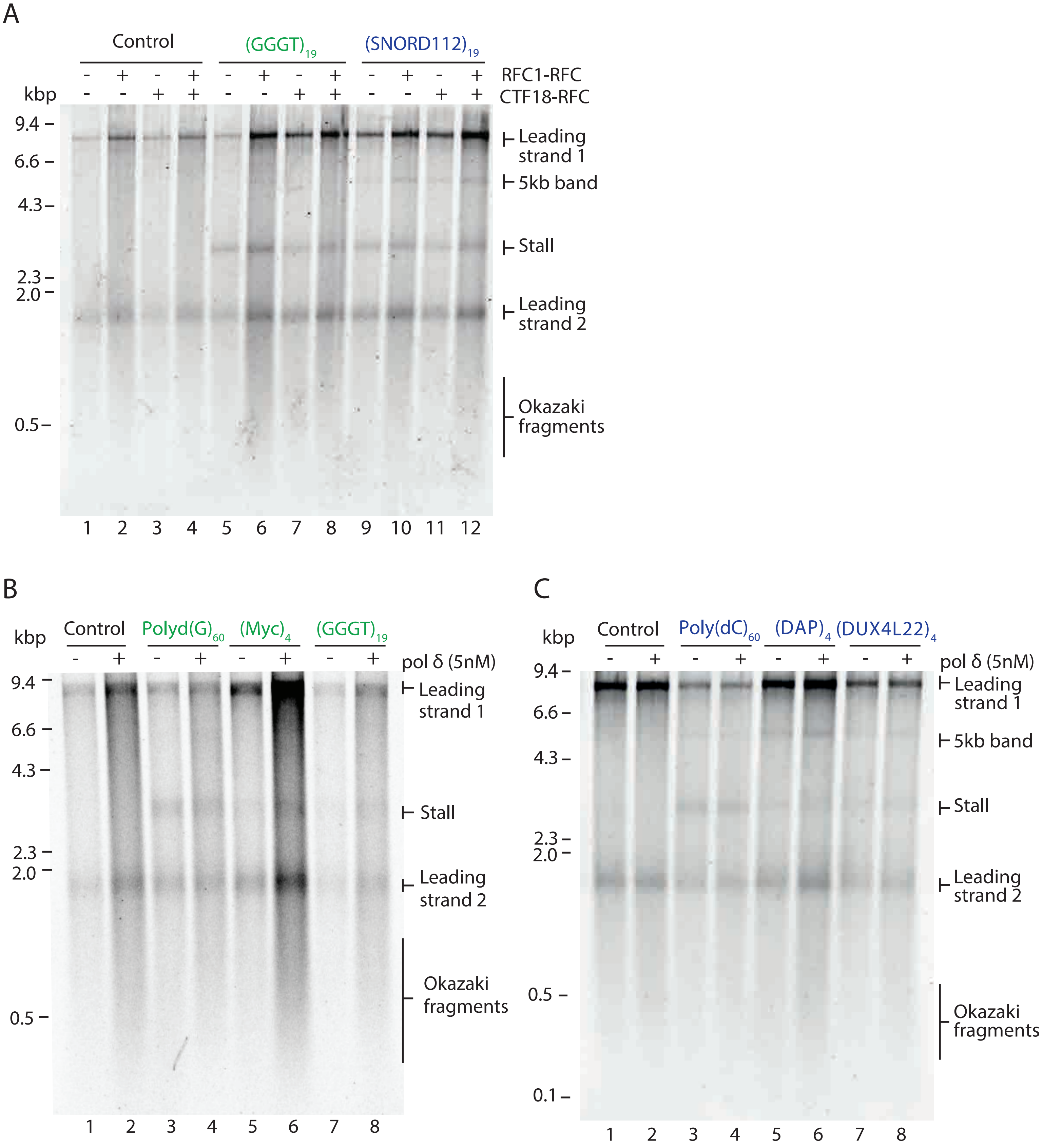


**Figure S8. Pol δ and CTF18-RFC do not affect replisome stalling at G4s or iMs.**
(A) Replication of substrates containing a G4 ($(GGGT)_{19}$) or iM ($(SNORD112)_{1}$) in the presence or absence of RFC1-RFC or CTF18-RFC as indicated. Products were visualised on a denaturing agarose gel.
(B and C) Analysis of replication products of G4 (B) or iM (C) substrates in the presence or absence of pol δ on denaturing agarose gels.

Figure S9

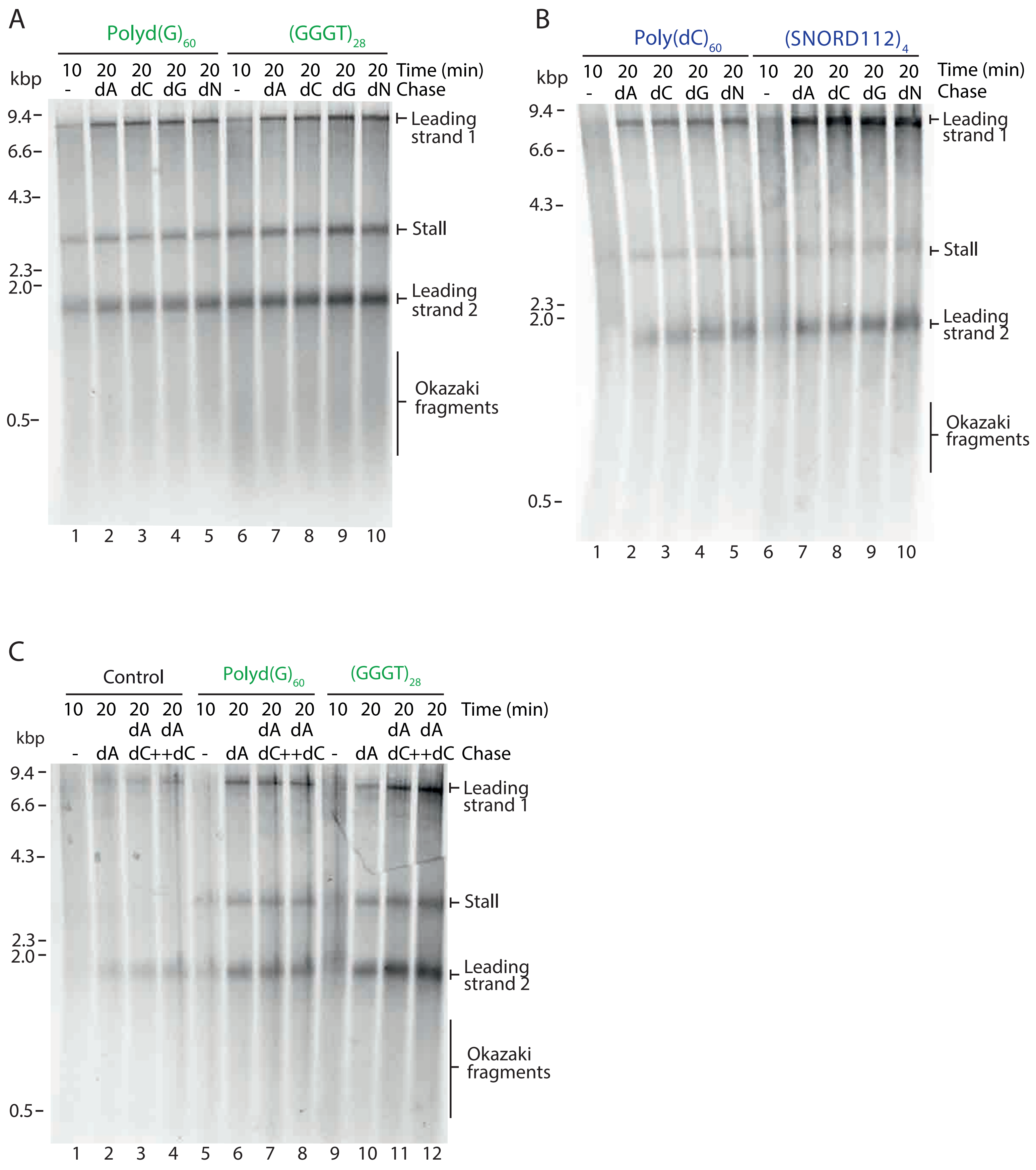


**Figure S9. Excess dNTPs can not rescue stalling at G4s or iMs.**
(A and B) Pulse-chase experiments carried out with the indicated templates.
Reactions were initiated with radiolabelled dATP for 10 min and chased for another 10 min with either excess 'cold' dATP alone (dA) or in combination with dCTP (dC), dGTP (dG), or all four dNTPs (dN). (C) As per A and B but reactions were chased with either 'cold' dATP alone or in combination with the same excess of dCTP or with double the excess of dCTP as dATP.

Figure S10

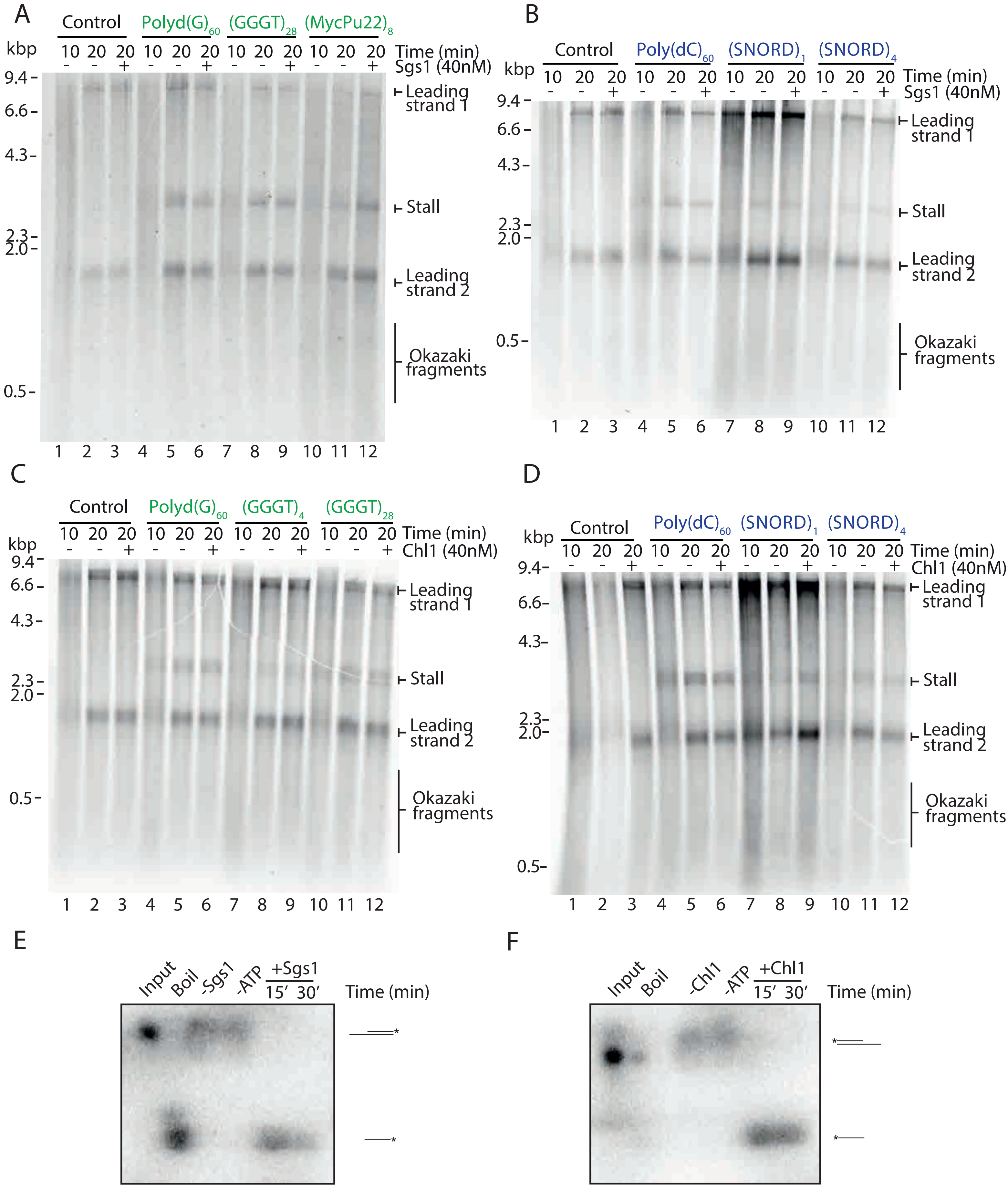


**Figure S10. Sgs1 and Chl1 are unable to rescue replication stalling at G4s or iMs.**
(A and B) Pulse-chase experiments carried out with the indicated templates. Reactions were initiated with radiolabelled dATP. After a 10 min pulse, either Sgs1 (A and B) or Chl1 (C and D) was added with the chase and samples taken after another 10 min. (E and F) Helicase unwinding assays using purified Sgs1 (E) and Chl1 (F). Time points were taken as indicated. Products were run on 10% TBE gels. Input and boiled substrates were used as controls to visualise where original and unwound substrates run on the gel. Assays demonstrate that both Sgs1 and Chl1 show robust unwinding activity.
